# Human DNA polymerase delta is a pentameric holoenzyme with dimeric p12 subunit: Implications in enzyme architecture and PCNA interaction

**DOI:** 10.1101/525485

**Authors:** Prashant Khandagale, Doureradjou Peroumal, Kodavati Manohar, Narottam Acharya

**Affiliations:** Laboratory of Genomic Instability and Diseases, Department of Infectious Disease Biology, Institute of Life Sciences, Bhubaneswar-751023, India

**Author notes:** Correspondence to: Narottam Acharya.

**Keywords:** Processivity, formaldehyde cross-linking, size exclusion chromatography, Cdm1, Replication foci, DNA replication

## Abstract

Human DNA polymerase delta (Polδ), a holoenzyme consisting of p125, p50, p68 and p12 subunits, plays an essential role in all the three DNA transaction processes. Herein, using multiple physicochemical and cellular approaches we found that the p12 protein forms a dimer in solution. *In vitro* reconstitution and pull-down of cellular Polδ by tagged p12 authenticates pentameric nature of this critical holoenzyme. Further, a consensus PIP motif at the extreme carboxyl terminal tail and a homodimerization domain at the amino-terminus of the p12 subunit were identified. Our mutational analyses of p12 subunit suggest that _3_RKR_5_ motif is critical for dimerization that facilitates p12 binding to IDCL of PCNA via its PIP motif _98_QCSLWHLY_105_. Additionally, we observed that oligomerization of the smallest subunit of Polδs is evolutionarily conserved as Cdm1 of *S. pombe* also dimerzes. Thus, we suggest that human Polδ is a pentameric complex with a dimeric p12 subunit; and discuss implications of p12 dimerization in regulating enzyme architecture and PCNA interaction during DNA replication.

## Introduction

Accurate and processive DNA synthesis by DNA polymerases (Pol) during chromosomal DNA replication is essential for the preservation of genomic integrity and for the suppression of mutagenesis and carcinogenesis (1). Eukaryotic chromosomal DNA replication is coordinated by three essential replicases: Polα, Polδ, and Polε (2,3). Based on biochemical and genetic studies mostly those carried out in the budding yeast, it has been proposed that Polα initiates DNA replication by synthesizing a short RNA-DNA primer, and is followed by loading of DNA clamp PCNA by its loader RFC. Polδ not only can synthesize “Okazaki fragments” in lagging strand, it can also take part in leading strand DNA synthesis (4,5). The precise role of Polε remains controversial but studies indicate that it plays a role in initiation and leading strand synthesis of DNA replication (3). The mechanism of DNA replication in higher eukaryotes is yet to be deciphered; however, the *in vitro* SV40 replication system suggests requirement of human Polα and Polδ to complete DNA synthesis (2,6). Irrespective of their different roles in DNA replication, these DNA polymerases possess certain commonalities like multisubunit composition, and the largest subunits only possessing signature sequences of a B-family DNA polymerase having catalytic activity (7). The accessory structural subunits of DNA polymerases do not show any sequence similarities between them, therefore understanding their functions is quite intriguing.

Among the replicative DNA polymerases, the subunit composition of Polδ varies from organism to organism. While ScPolδ consists of three subunits Pol3, Pol31 and Pol32, Polδ from *S. pombe* possesses four subunits Pol3, Cdc1, Cdc27 and Cdm1 (5,8–10). The mammalian Polδ holoenzyme consists of p125 as the catalytic subunit, the yeast homologue of Pol3; p50, p68 and p12 structural subunits (11). The accessory subunits p50 and p68 are the equivalents of Pol31/Cdc1 and Pol32/Cdc27 subunits, respectively. The p50/Pol31/Cdc1 subunit makes a connecting bridge between the catalytic subunit p125/Pol3 and p68/Pol32/Cdc27, and is indispensable for cell viability. Even though Pol32 is not essential for cell survival in *S. cerevisiae,* in its absence cells exhibit sensitivity to both high and cold temperatures, and susceptibility to genotoxic stress (12). Contrarily, deletion of Cdc27 in *S. pombe* is not achievable (13). The non-essential p12 subunit is the Cdm1 homologue and is apparently absent in *S. cerevisiae.* Yeast two hybrid and co-immunoprecipitation analyses suggested a dual interaction of p12 with p125 and p50; however, the modes of binding are yet to be defined (14). *In vitro* reconstitution has facilitated purification of four different subassemblies of human Polδ such as p125 alone, p125-p50 (core complex), p125-p50-p68 and p125-p50-p68-p12 complexes for biochemical studies. Reports also suggest that the subunit composition of human Polδ may alter *in vivo* with cellular response to genotoxic burden (15,16). Upon treatment of human cells with genotoxins such as UV, methyl methanesulfonate, HU and aphidicolin, the p12 subunit undergoes rapid degradation to result in a trimeric human Polδ (p125/p50/p68) equivalent to yeast form with higher proof reading activity (17). Thus, p12 subunit seems to play a crucial role in regulating Polδ function.

The function of Polδ as a processive DNA polymerase mostly depends upon its association with PCNA that acts as a sliding clamp (18). The interaction of PCNA-binding proteins with PCNA is mediated by a conserved PCNA-interacting protein motif (PIP-box) with a consensus sequence Q-x-x-(M/L/I)-x-x-FF-(YY/LY). Previously, we have shown that all the three subunits of ScPolδ functionally interact with trimeric PCNA mediated by their PIP motifs (5). To achieve higher processivity *in vitro*, all the three PIP boxes are required; however for cellular function of ScPolδ, along with PIP of ScPol32, one more PIP box either from Pol3 or Pol31 subunit are essential. Similarly, reports from human Polδ studies suggest that all the four subunits of hPolδ are involved in a multivalent interaction with PCNA and each of them regulates processive DNA synthesis by Polδ (19). Except in p68 and p50, PIP motifs in p125 and p12 have not been mapped (20,21). Studies based on far-Western and immunoprecipitation analyses revealed that the first 19 amino acids of p12 are involved in PCNA interaction, although the region lacks a canonical PIP box sequences (14,22).

Therefore, in this study we have re-investigated the interaction of p12 with other Polδ subunits and PCNA. Our results indicate that the smallest subunit p12 exists as a dimer in solution to establish a dual interaction with both p125 and p50 subunits of Polδ. The dimerization motif has been mapped to the amino-terminal end and a novel conserved PIP box has been identified in the c-terminal tail of p12. Importantly, dimerization of p12 regulates its interaction with PCNA. Based on our observations we propose that human Polδ exists in a pentameric form in the cell in addition to other subassemblies; and the effect of p12 dimerization on PCNA binding with various subassemblies have been discussed.

## Results

### Oligomerization of p12 subunit of hPolδ

In **s**everal studies, human Polδ holoenzyme has been purified either from insect cell line or bacterial systems by using standard chromatography techniques for biochemical characterizations (11,23,24). Although most of the purified Polδ used in various enzymatic assays was containing all the four subunits p125, p50, p68 and p12; in most cases the proposed stochiometry of each subunit in the holoenzyme 1:1:1:1 was less convincing. Especially the smallest subunit p12 was more than 1:1 ratio with respect to others. The band intensity of p12 was always either similar or more than that of other subunits (23,24). Even when we attempted to purify hPolδ holoenzyme by using GST-affinity beads, the band intensity of p12 protein was consistently higher than the other subunits (Supplementary Fig. 1). Such discrepancy in Polδ composition could arise due to either oligomeric status of p12 or multi-subunit p12 interaction with p125 and p50 or due to staining artifacts. To examine potential oligomerization status of p12 subunit, yeast two hybrid assay and native PAGE analysis were carried out (Fig. 1). p12 orf was fused in frame with both *GAL4* activation (AD), and *GAL4* binding domains (BD). Other hPolδ subunit orfs were fused with the *GAL4* binding domain only. The HFY7C yeast reporter strain harboring the AD-p12 plasmid was co-transformed with any of the BD-Polδ subunit plasmids, and selected on Leu^−^Trp^−^ SDA plate. The interactions of p12 with any subunits of Polδ in these transformants were analyzed by selecting them on plate lacking histidine. Growth on His^−^ plate demonstrates interaction between the two fusion proteins as only the binding of two proteins make it possible to form an intact *GAL4* activator to confer *HIS* expression. As reported earlier, p12 interaction was observed with p125 and p50 but not with p68 (Fig. 1 A, sectors 1, 2, and 3) (14). Surprisingly, p12 subunit also interacted with itself to give *HIS* expression (sector 4); while no growth was observed in the negative controls (sector 5 and 6). As no growth was observed to show p12 and p68 binding, our results suggest that p12 makes specific multivalent interaction with itself, p125 and p50 subunits; and such interactions are not mediated through any yeast proteins.

**Figure 1:**
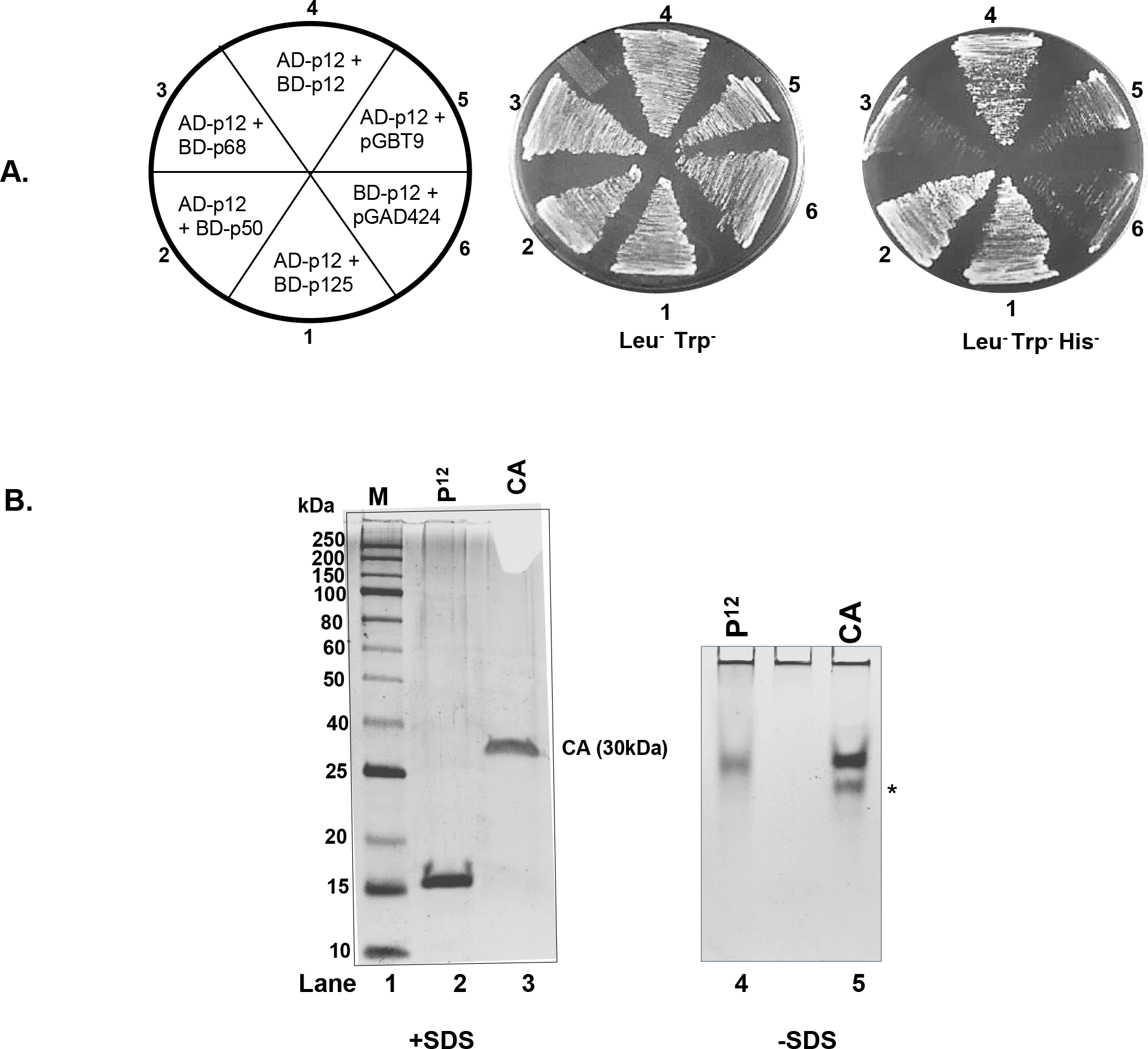
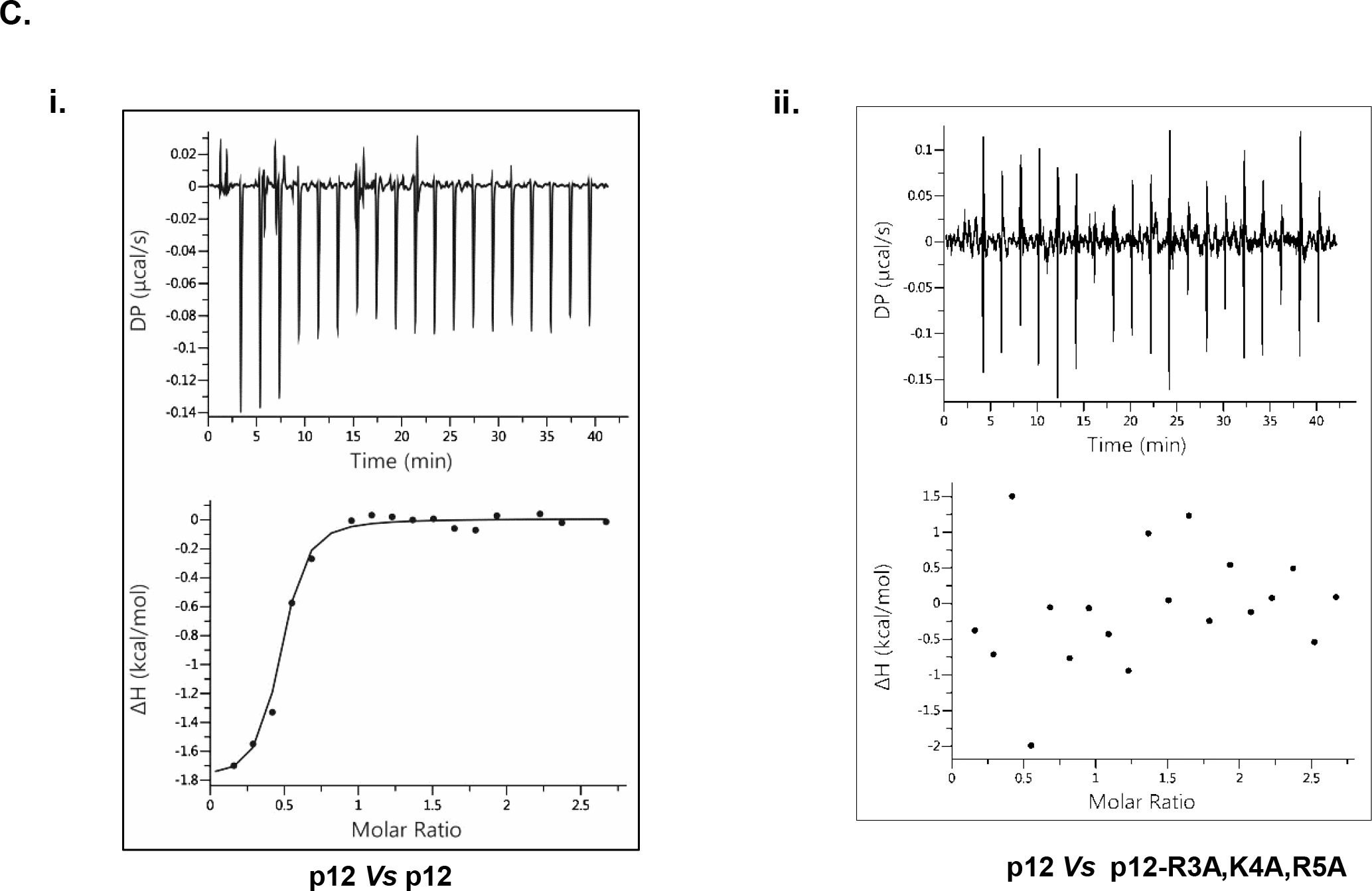
Interaction of p12 with hPolδ subunits. **A**. Yeast two hybrid analysis showing interaction of p12 with various subunits of hPolδ. HFY7C yeast transformants with various GAL4-AD and BD fusions were selected on SD media plates lacking leucine, tryptophan with and without histidine amino acids. Sector 1, AD-p12 + BD-p125; Sector 2, AD-p12 + BD-p50; Sector 3, AD-p12 + BD-p68; Sector 4, AD-p12 + BD-p12; Sector 5, AD-p12 + pGBT9; and Sector 6, AD-p12 + pGAD424. **B**. Purified p12 protein was resolved in native and SDS containing-PAGEs. Lane 1, MW; Lane 2 and 4, p12; Lane 3 and 5, Carbonic anhydrase. **C**. ITC analysis of p12 to wild type (i) or RKR-mutant p12 (ii) at 25°C. In each panel, the upper half shows the measured heat exchanges during each protein injection. The lower half of each panel shows the enthalpic changes as a function of the molar ratio of p12 to wild type or RKR mutant p12 monomer. Circles and lines denote the raw measurements and the fitting to a one set of identical sites.

Further to ascertain oligomerization of p12, protein was purified to near homogeneity from bacterial cells by using GST-affinity column chromatography, GST-tag was cleaved off by PreScission protease, and analyzed by both native and SDS containing polyacrylamide gel electrophoresis (PAGE). The predicted MW of p12 is ~12 kDa with a pI of 6.3. p12 is known to possess abnormal migration in SDS-PAGE (25) and similarly we observed the protein to migrate at ~15 kDa molecular weight size position (Fig. 1 B lane 2). By taking advantage of native PAGE analysis where proteins resolve based on their charge and hydrodynamic size, we found that p12 migrated at similar position with Carbonic anhydrase (CA) which is a protein of 30 kDa with pI 6.4 (lane 4 and 5). Thus, the co-migration of two proteins indicate that both CA and p12 possess similar mass to charge ratio and the slower migrating p12 is potentially a dimeric complex.

By taking advantage of Isothermal calorimetry (ITC) technique, the oligomerization of p12 protein was examined (Fig. 1C i). p12 was placed both in the sample cell and in the syringe of the calorimeter and binding was analyzed by monitoring the change in entropy. The ΔH was ~ −1.82 kcal/mol, the ΔG was − 9.48 kcal/mol, and the *Kd* for the complex was ~146 nM. The number of ligand binding site as derived from the ITC analysis was found to be ~0.6, which is closed to 1:1 binding of p12 monomers. Thus, our both *in vivo* and physiochemical study clearly indicate homodimerization of p12 subunit.

### Cellular existence of oligomeric p12 in Polδ complex and co-localization with PCNA

In order to decipher p12 oligomerization in its cellular state, Polδ holoenzyme was immunoprecipitated from the cell lysate transfected with GFP-p12 by using either anti-GFP (i) or anti-p125 (ii) antibody (Fig. 2 A). Irrespective of any antibody used for the pull-down assay, we detected presence of five subunits of Polδ in the beads. Each of the four native subunits (p125, p68, p50 and p12) were detected by probing with subunit specific antibody where as GFP-p12 was detected by anti-GFP antibody. While GFP-p12 pulled down cellular heterotetrameric Polδ complex, anti-p125 antibody precipitated Polδ complex with both the forms of p12 (native and GFP-tagged). Thus, even in the cellular context, p12 exists as an oligomer in the Polδ complex.

**Figure 2:**
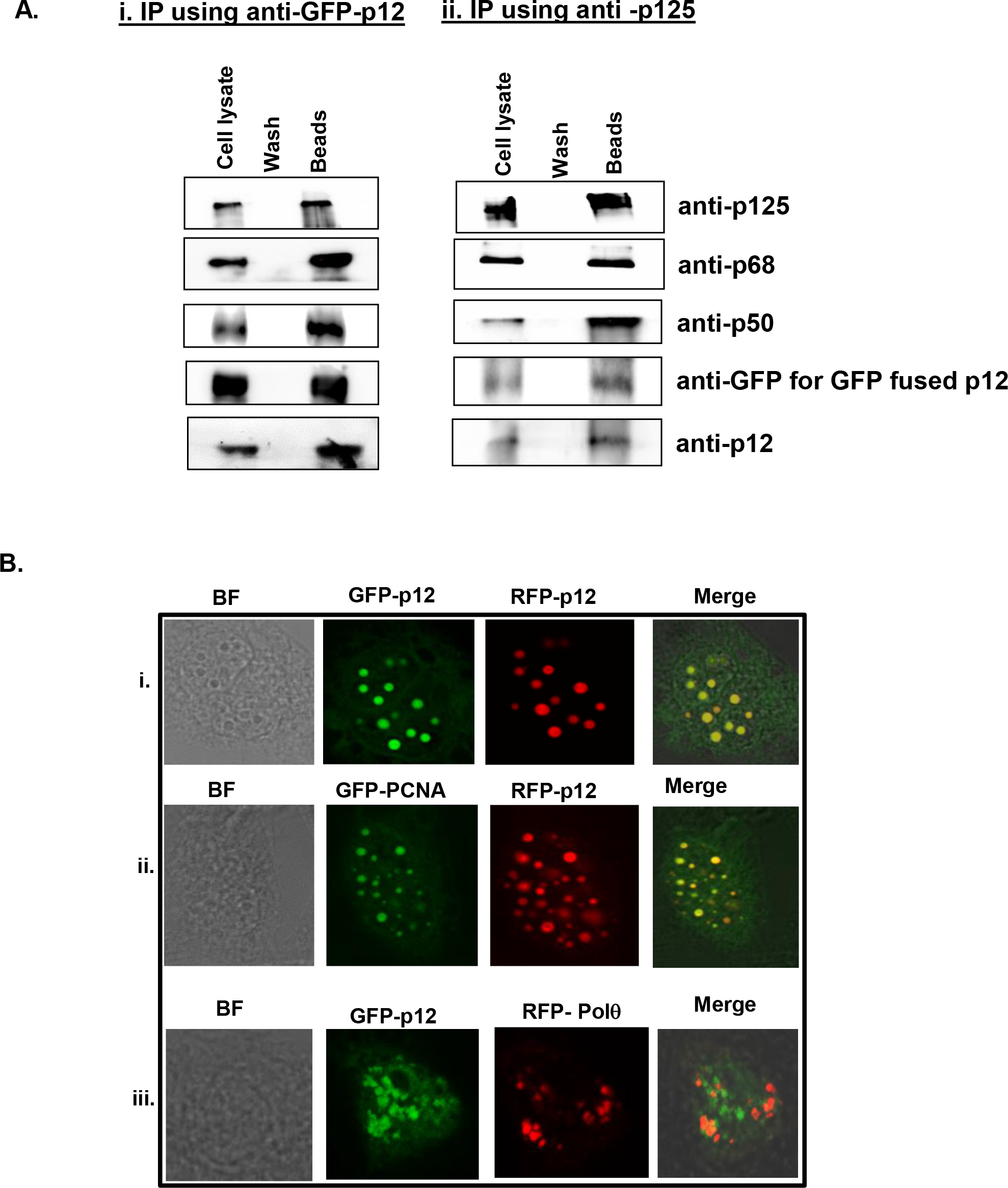
Existence of p12 dimers in the cellular context of Polδ. **A.** Native human Polδ was co-immuno-precipitated from HEK293 cells transfected with GFP-p12. Either GFP-p12 (i) or p125 (ii) was used to immunoprecipitate cellular Polδ. After through washings, eluate was separated in 12 % SDS-PAGE, and presence of various subunits of hPolδ was detected by subunit specific antibody. GFP-p12 was detected by anti-GFP antibody. **B.** Nuclear co-localization of p12 and PCNA. CHO cells were co-transfected with GFP-p12 and RFP-p12 (i) or GFP-PCNA and RFP-p12 (ii) or GFP-p12and RFP-Polθ(iii).After 48 hrs, cell were fixed and mounted as described in experimental procedures section and images were taken using Leica TCS SP5 at 63 X objective.

PCNA that functions as a cofactor for DNA polymerases orchestrates the replisome by recruiting multiple proteins involved in DNA transaction processes. Using microscopy, it was shown that PCNA forms distinct foci or replication factories indicative of active DNA replication entities within the nucleus (26). The p68 and p50 subunits of hPolδ were deciphered to form foci and co-localized with PCNA in several cell lines (27,28). To examine physiological relevance of p12 oligomerization and PCNA interaction, GFP-PCNA or GFP-p12 with RFP-p12 fusion constructs were transfected to CHO cell line. As shown in Fig.2 B, irrespective of GFP (stained green) or RFP (stained red) fusion, p12 formed discrete compact foci and the patterns were very similar to reported p50 and p68 foci. Subsequent merging of foci in co-transfectant of GFP-p12 and RFP-p12 resulted in appearance of yellow foci (i). Thus, 100% coincidental accumulation of both the p12 foci suggests that both the proteins are part of replication unit and function together. Similarly, we have also observed sub-cellular co-localization of p12 with PCNA as yellow foci appeared by merging foci of GFP-PCNA and RFP-p12 (ii). However, in similar condition, we did not notice any co-localization of GFP-p12 and RFP-Polθ foci (iii). We suggest from these observations that p12 could function in replication factories as an oligomeric protein with PCNA.

### Identification of motifs involved in p12 dimerization and interaction with PCNA

To identify dimerization and PIP motifs in p12, primary sequences from human, mouse, bovine and *S. pombe* were aligned (Fig. 3 A). The CLUSTAL W alignment analysis showed a high degree of amino acid conservation of p12 sequences in mammals, and they showed only ~17% identity with *S. pombe* orthologue Cdm1. The carboxyl termini of these proteins displayed better conservation than the amino termini. Cdm1 is composed of 160 aa and the divergence is apparently due to possession of an insert of about 38 amino acids exactly in the middle of the p12 homology sequences in Cdm1. We thought to examine the role of two highly conserved sequences located at the extreme ends of p12: a basic tripeptide sequence _3_RKR_5_ (henceforth referred as RKR motif) motif and a putative PIP box sequence _98_QCSLWHLY_105_ (henceforth referred as PIP motif). The putative PIP motif is located in the carboxyl terminal tail of p12, which is the usual position of PIP in most of the DNA polymerases and appears to be very similar to known PIP sequences (Fig. 3 B). Since, an earlier study reported _4_KRLITDSY_11_ (a part of RKR motif) as a PCNA binding region of p12 (14), we wanted to compare the model structure of this peptide with _98_QCSLWHLY_105_. Structurally PIP box sequences are highly conserved and formation of a 3_10_ helix is a characteristic of such sequences. The amino acid stretch encompassing RKR (1-MGRKRLITDSYPVK-14) and PIP (92-GDPRFQCSLWHLYPL-106) domains were used for peptide structure prediction by using PEP-FOLD3 server (http://bioserv.rpbs.univ-paris-diderot.fr/services/PEP-FOLD3/) rather than using a known template based prediction to avoid any biasness. Further the models were validated by SAVES and Ramachandran plot (Supplementary Fig. 2 A and B), which showed most of the residues in allowed regions. Our structural prediction suggested first 10 amino acids of RKR motif to form a α-helix whereas the PIP motif of p12 forms a typical 310 helix, the structure that snugly fits into the IDCL domain of PCNA. The p12 PIP structure was further aligned with available X-ray crystal structures of PIP peptide from p21 (1AXC) and p68 (1U76). The superimposition shows a high degree of similarity between the PIP motifs (Fig. 4 C). Similarly, p12 peptide structure was aligned with that of p68 PIP-hPCNA co-crystal structure, and a remarkable overlapping between the structures was observed (Fig. 3 C). Thus, our *in silico* analysis indicated that the C-terminal PIP sequence _98_QCSLWHLY_105_ is the most probable motif to interact with PCNA than the N-terminal RKR motif _4_KRLITDSY_11_.

**Figure 3:**
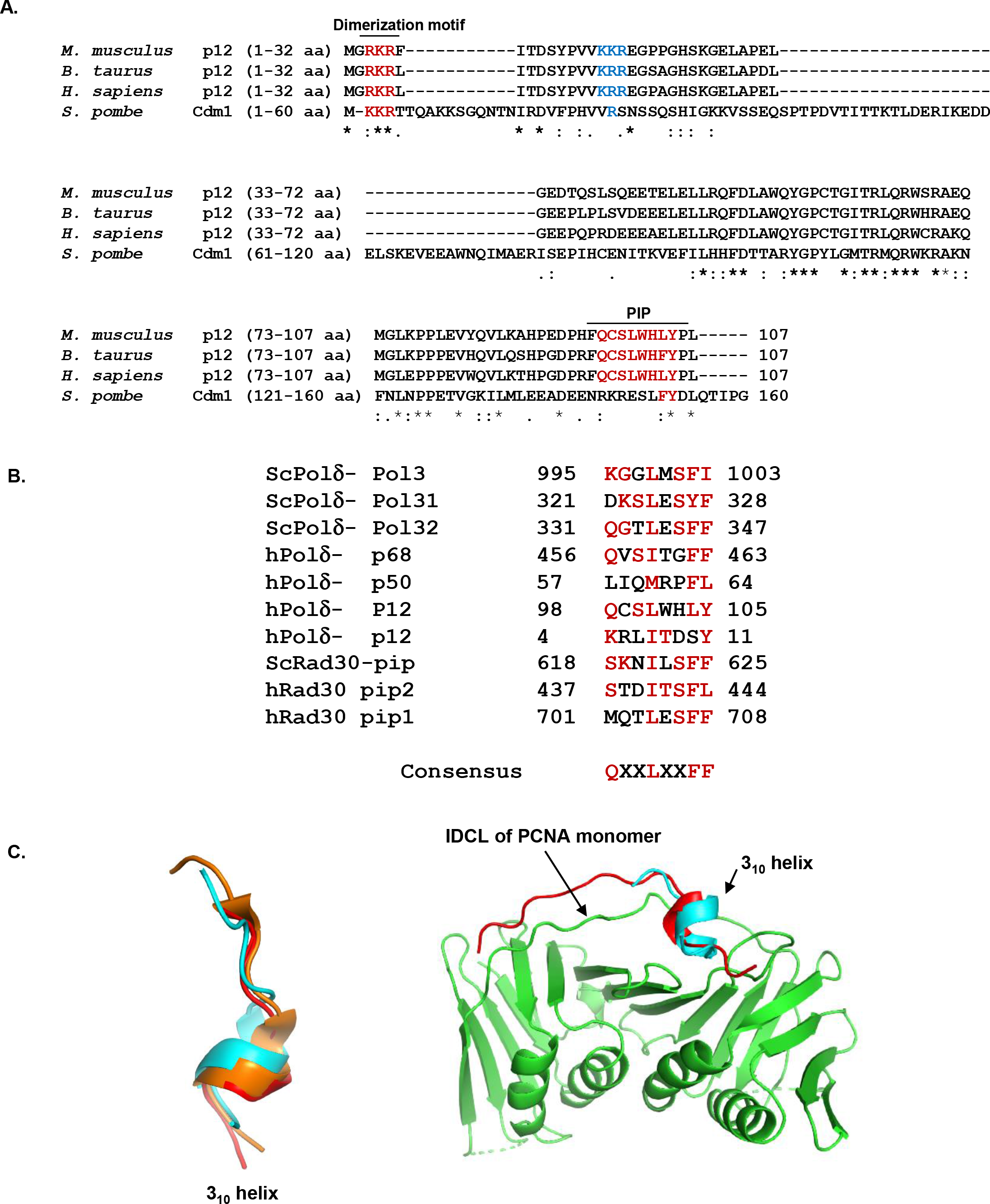
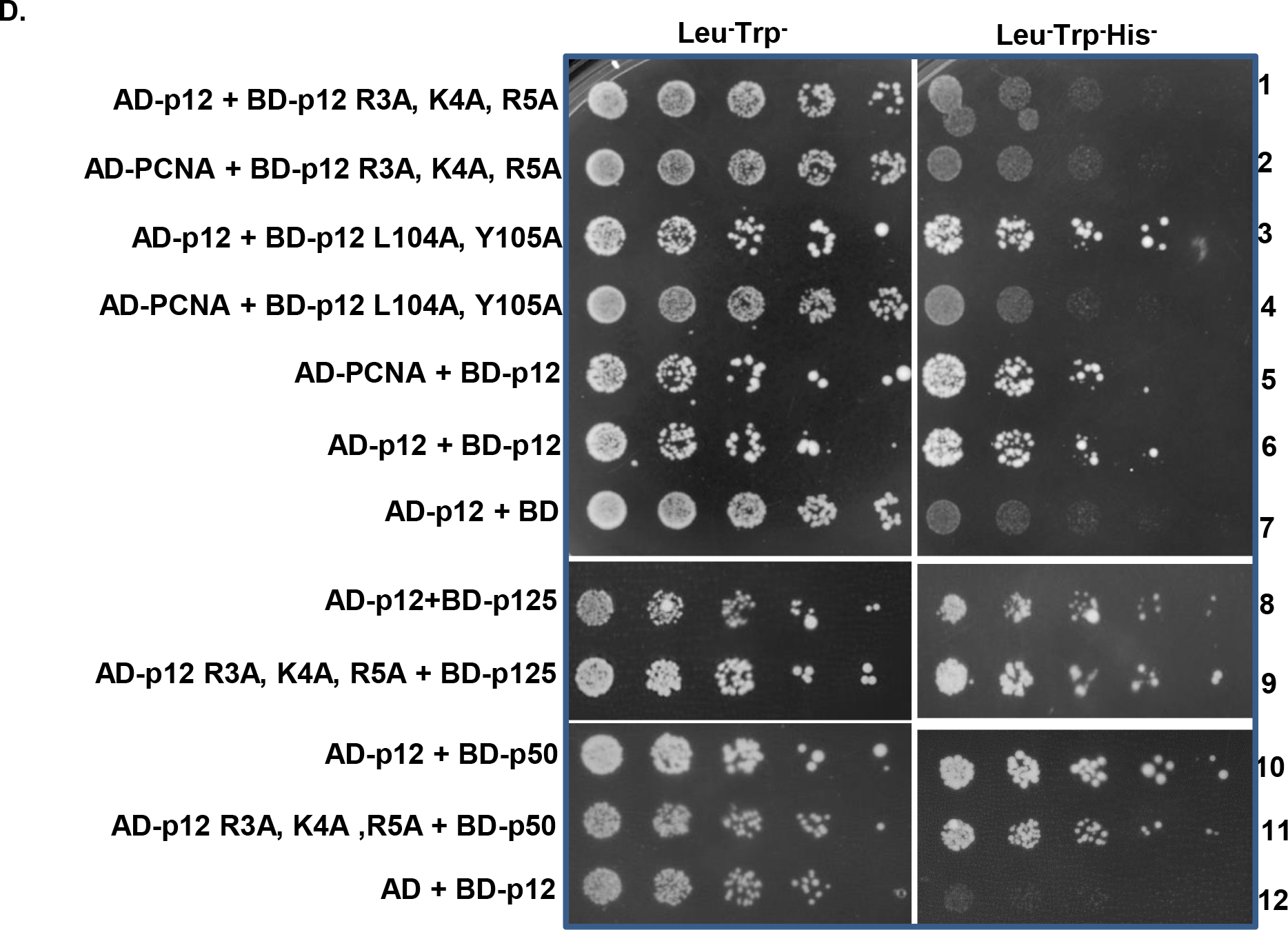
Identification of p12 motifs involved in dimerization and PCNA interaction. **A.** The primary sequences of the 4^th^ subunit of Polδs from human (*Homo sapiens*), mouse (*Mus musculus*), bovine (*Bos Taurus*) and fission yeast (*Schizosaccharomyces pombe*) were aligned by CLUSTALW. Conserved dimerization motif and PIP-box sequences are colored brown. Secondary structure prediction showed coiled structures with three putative α helices. Identical residues are marked with*, conserved residues are marked with and less conserved residues are marked with •. **B.** Identified PIP motif residues from DNA polymeraseη and δ subunits were aligned and compared that with putative PIP sequences in p12. Repeated residues in more than one sequences are highlighted. **C**. Superimposition of model 310 helix structure of p12 PIP (cyan) with available PIP structures from p21 (orange) and p68 (red) peptides without and with human PCNA monomer. **D.** Yeast two hybrid analysis showing interaction of mutants p12 with wild type p12 and PCNA. HFY7C yeast transformants with various GAL4-AD and BD fusions were selected on SD media plates lacking leucine, tryptophan with and without histidine amino acids. Row 1, AD-p12 + BD-p12 R3A, K4A, R5A; Row 2, AD-PCNA + BD-p12 R3A, K4A, R5A; Row 3, AD-p12 + BD-p12 L104A, Y105A; Row 4, AD-PCNA + BD-p12 L104A, Y105A; Row 5, AD-PCNA + BD-p12; Row 6, AD-p12 + BD-p12; Row 7, AD-p12 + pGBT9. Row 8, AD-p12 + BD-p125; Row 9, AD-p12 R3A, K4A, R5A + BD-p125; Row 10, AD-p12 + BD-p50; Row 11, AD-p12 R3A, K4A, R5A + BD-p50; and Row 12, AD-p12 + pGBT9.

**Figure 4:**
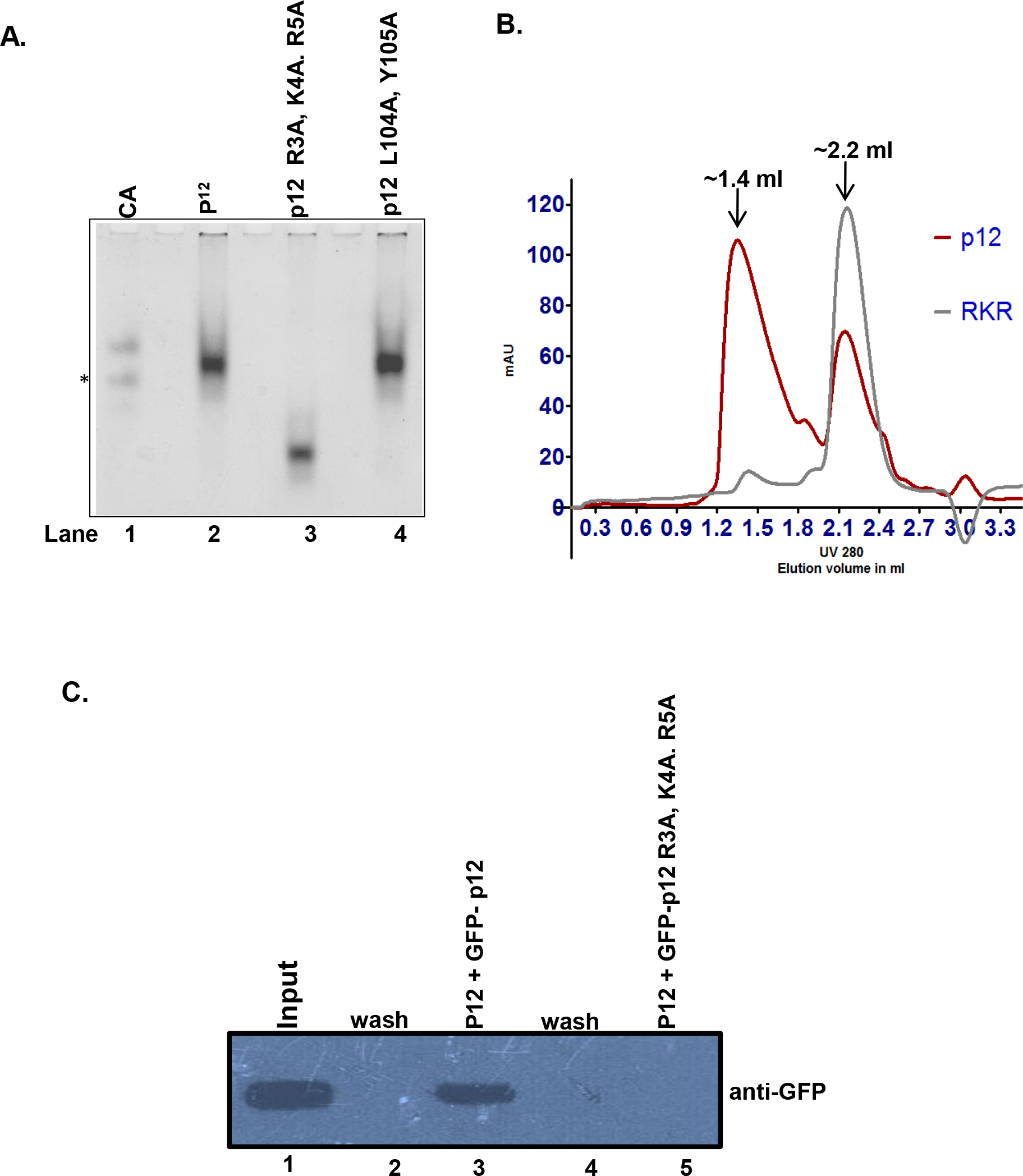
RKR-motif is involved in dimerization. **A.** R3A, K4A, R5A and L104A, Y105A p12 proteins were resolved in native PAGE. Lane 1, CA; Lane 2, p12; Lane 3, R3A, K4A, R5A; and Lane 4, L104A, Y105A. * represents degraded CA. **B.** About 10 μg of wild type (red line) and R3A, K4A, R5A (grey line) p12 proteins were subjected to size exclusion chromatography. Two elution peaks at ~1.4 ml and ~2.2 ml were observed representing a dimer and monomer populations, respectively. **C.** Immunoprecipitation of GFP-p12 by FLAG-p12 (lane 3) but not of GFP-R3A, K4A, R5A mutant (lane 5). Lane 1, cell lysate input; Lanes 2 and 4 are washings from beads.

To provide experimental evidence to our *in silico* prediction, two p12 mutants were generated by mutating R3, K4, R5 and L104, Y105 to alanines; and their binding to wild type p12 and PCNA were analyzed by yeast two hybrid approach (Fig. 3 D). As depicted, while transformants of BD-p12 with AD-p12 or AD-PCNA grew on SDA plate lacking leucine, tryptophan and histidine amino acids (rows 5 and 6); both R3A, K4A, R5A and L104A, Y105A p12 mutants failed interact with PCNA and thus no growth was observed (rows 2 and 4) similar to the vector control (row 7). Interestingly, R3A, K4A, R5A mutant is also defective in p12 interaction in yeast cells but not the L104A, Y105A mutant (compare row 1 with 3). These *in vivo* results suggest that while RKR-motif plays a dual role in dimerization and PCNA interaction, _98_QCSLWHLY_105_ is only involved in PCNA binding.

Further, we wanted to examine whether dimerization of p12 is also required for other Polδ subunit interactions. Yeast two hybrid analyses of co-transformants harboring AD-R3A, K4A, R5A with BD-p125 or BD-p50 demonstrated that mutation in this motif has no effect on Polδ’s subunit interaction as transformants grew efficiently on Leu-Trp-His-plate (Fig. 3 D, sectors 8-12). Thus, dimerization of p12 is not required for p125 or p50 binding. However, it is not clear whether p125 and p50 bind to the same or different regions in p12. Analyses of various deletion constructs in p12 did not reveal any significant results to suggest any interaction domain (data not shown). Nonetheless, dimerization will facilitate binding of p125 and p50 subunits of Polδ to separate monomer of p12 dimer.

### RKR-motif of p12 is critical for dimerization

Multiple basic amino acid motifs such as RKR/KKR/KRK in other proteins are known to play crucial roles in a variety of cellular processes such as their retention and exit from ER, nuclear localization as well as the gating of K+ channels (29–32). Such motifs are also found to be involved in protein-protein interaction (33–35). To ensure the involvement of RKR-motif in p12-p12 dimerization, mutant proteins R3A, K4A, R5A and L104A, Y105A were purified to near homogeneity and analyzed in SDS-PAGE (Supplementary Fig. 3 lanes 3 and 4). The mutant proteins were co-migrated with the wild type p12 close to the dye front and thus mutations in these residues had no obvious effect on protein mobility and stability. When the proteins were further resolved in non-denaturating PAGE, both wild type and L104A, Y105A proteins were co-migrated; and their migration was similar to CA as shown earlier (Fig. 4 A, compare lanes 1 with 2 and 4). However, R3A, K4A, R5A mutant p12 was migrating much faster than the other two p12 proteins in native gel as a monomer (Fig. 4 B compare lane 3 with lanes 2 and 4). Therefore, we concluded that RKR motif is absolutely required for dimerization, and mutations in _98_QCSLWHLY_105_ motif have no effect on such a function of p12.

To rule out the possibility that the faster migration of RKR mutant in non-denaturating gel is not due to because of change in residual charge in the protein rather due to abolition of dimerization, we compared the gel filtration elution profiles of wild type and R3A, K4A, R5A mutant p12 by separating equal amount of proteins through S_200_ molecular exclusion chromatography at physiological salt concentration. While the wild type p12 protein eluted in two peaks of volume at ~1.4 ml and ~2.2 ml corresponding to an oligomeric and monomeric status of the protein (red line), R3A, K4A, R5A mutant only eluted (grey line) later at a single peak volume of ~2.2 ml (Fig. 4 B). This also rules out any change in stokes radius of p12 proteins due to mutation in the dimerization domain as a portion of wild type co-elutes with mutant p12.

Co-immunoprecipitation experiment was also carried out in the cell lysates harboring FLAG-p12 and GFP-p12 or GFP-p12 R3A, K4A, R5A by using anti-FLAG antibody conjugated beads; and further probed with anti-GFP antibody (Fig. 4 C). While anti-FLAG antibody could pull down wild type p12 as detected by anti-GFP antibody, it did not precipitate the RKR mutant (compare lane 3 and 5). Corroborating with our yeast two hybrid results, native PAGE, size exclusion chromatography and pull-down assays clearly demonstrate that indeed RKR-motif is a protein-protein interaction domain and is essential for p12 dimerization.

### Formaldehyde cross-linking reveals dimerization of p12

Our native PAGE and ITC analyses suggested potential dimeric nature of p12 as it was migrating at similar position with CA (30 kDa). To estimate the exact number of p12 molecules in the oligomeric complex, we employed formaldehyde cross-linking assay. Reagents like formaldehyde or glutaraldehyde cross-links neighboring lysine or arginine residues of proteins to form a stable complex that can even withstand SDS (36,37); moreover, our mutational analysis deciphered the involvement of RKR motif in dimerization. Therefore, purified recombinant proteins were cross-linked with formaldehyde, analyzed on 12 % SDS-PAGE and detected by Coomassie brilliant blue staining (Fig. 5 A). Upon treatment with the cross-linker, both wild type and L104A, Y105A mutant p12 proteins showed a concentration dependent cross-linked dimers (lanes 2, 3, 4, 10, 11 and 12) those were migrating below 32 kDa position and did not form any higher order oligomers, whereas R3A, K4A, R5A mutant protein remained as monomer (lanes 6, 7 and 8). Without any cross linker, all the three proteins migrated at the bottom of the gel (lanes 5, 9 and 13). In addition to _3_RKR_5_ motif, p12 also possess yet another similar multi-basic motif _15_KKR_17_ in only mammalian p12 (Fig. 3 A colored in blue). As R3A, K4A, R5A p12 mutant failed to form any oligomer in the presence of formaldehyde, we suggest that 15KKR17 sequences has no role in such a function and dimerization is specifically mediated by R_3_K_4_R_5_ motif.

**Figure 5:**
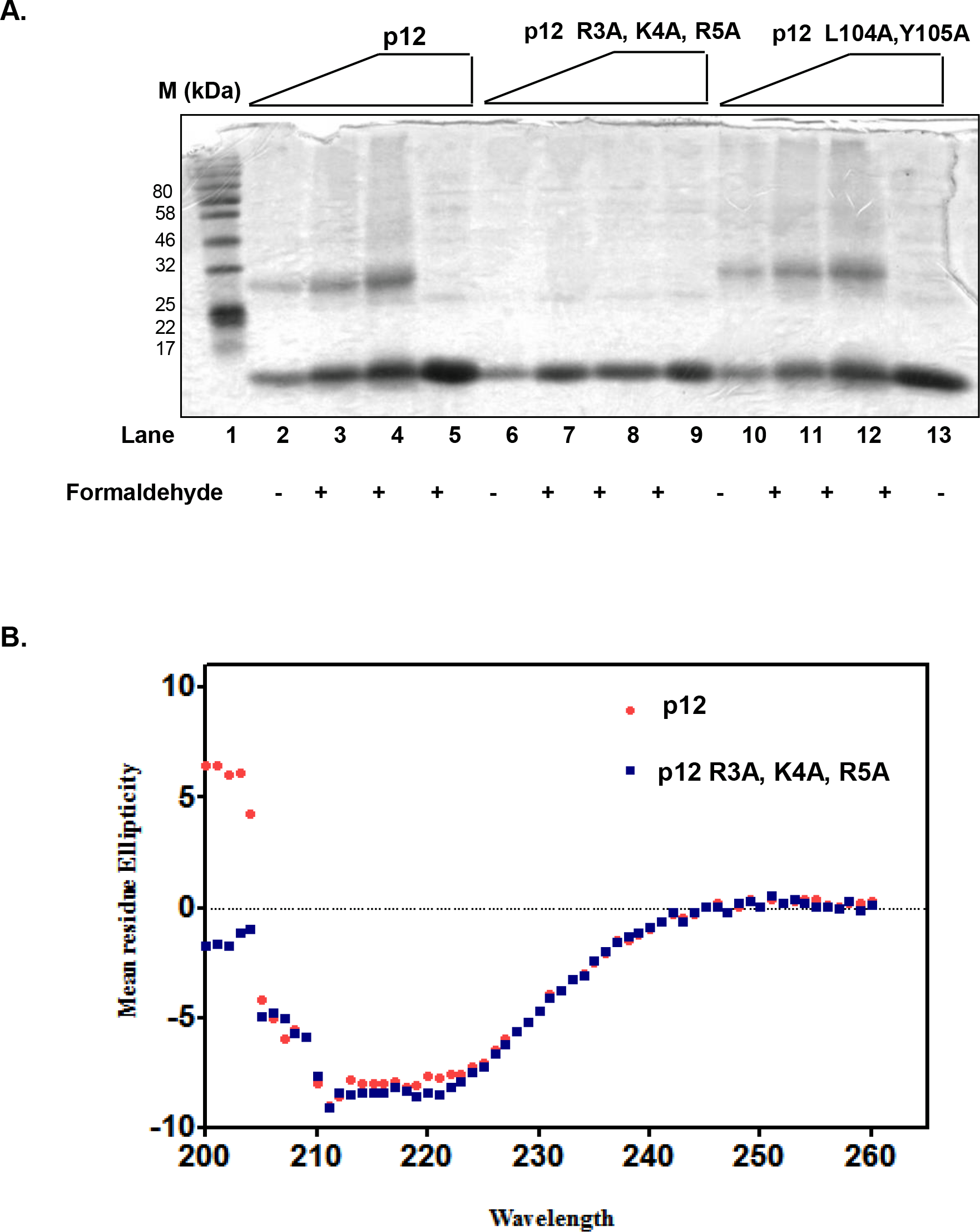
Formaldehyde cross-linking of p12 proteins. **A.** About 1-5 μg of p12 proteins were cross linked with 0.5% formaldehyde solution for 30 min at 25°C. After termination with SDS sample buffer, they were resolved in a 12 % SDS-PAGE. Lane 1, MW; Lanes 2-5, p12; Lanes 6-9, R3A, K4A, R5A and Lanes 10-13, L104A,Y105A. Lanes 5, 9 and 13, proteins treated similarly but without formaldehyde. **B**. Far UV-CD spectra of wild-type (red) and R3A, K4A, R5A mutant (blue) p12. CD spectra at pH 7.5 between 200 nm and 260 nm were recorded. Data represent values determined after solvent correction and after averaging each set (n=3).

Since, we did not detect dimerization of RKR-motif mutant in any of our assay, we wanted to rule out the possibility that this effect resulted from a significant change in p12 conformation. For this reason, we compared the CD spectra of the wild-type and R3A, K4A, R5A mutant p12 proteins (Fig. 5 B). The CD spectra determined in the “far-UV” region (200 to 260 nM) showed p12 to be enriched in α-structure as evident from the characteristic negative peaks at 208 nm and 222 nm. It also indicates that the mutant protein retains similar level of secondary structures as that of wild-type protein, which would suggest that the mutations do not cause a major perturbation of p12 structure. Even the ITC assays failed to detect any binding between wild type p12 and R3A, K4A, R5A p12 mutant (Fig. 1 C ii and D ii). Considering all these evidences, we conclude that p12 forms a dimer solely mediated by R_3_K_4_R_5_ motif.

### *In vitro* reconstitution of pentameric human Polδ holoenzyme

Various subassemblies of human Polδ such as p125 alone, p125-p50, p125-p50-p68, and p125-p50-p68-p12 have been purified *in vitro* by mixing various combination of purified proteins (11,38). In this study, Polδ holoenzyme was expressed by co-transforming two bacterial expression constructs GST-p125 and pCOLA234 (p50-p68-His-Flag-p12); and complex was purified to near homogeneity as described in the methods section. Taking advantage of strategically located PreScision protease site, cleaved Polδ complex was obtained in which only p12 subunit was amino-terminally FLAG tagged. To conclusively show two different forms of p12 in the holoenzyme, untagged p12 protein purified from bacterial GST-p12 system was added and the mixture was further incubated at 4 °C for 4 hrs. We argued that if p12 forms a dimer in the Polδ complex, untagged p12 will compete out some of the resident Flag p12 and a Polδ with five subunits will be appeared. Thus, the mixture was loaded into S_200_ minicolumn for separation and fractions were collected in a 96 well plate. Since we could not detect enough protein by coomassie staining despite our repeated trails, various fractions were analyzed in SDS-PAGE followed by detection of various subunits by probing with specific antibody (Fig. 6). To detect the presence of enriched holoenzyme fractions, the membrane was first probed with anti-p68 antibody (A) and further selected fractions were again separated in SDS-PAGE (B). As depicted in the figure, initial fractions were enriched in Polδ holoenzyme (A8-B2) and a clear shift of untagged p12 proteins was also detected. Interestingly, early fractions such as A8-A10 only showed majority of Flag-p12 and very little amount of untagged p12; whereas the later fractions convincingly showed presence of both tagged and untagged p12 subunits (A12-B2); it also suggests that early eluate are also pentameric complex. In this approach, we could purify two populations of Polδ5; (a) early Polδ5 fractions containing dimeric FLAG-p12 (A8-A10) and (b) later Polδ5 fractions where some of the FLAG-p12 were replaced with untagged p12 (A12-B2). This analysis clearly demonstrates that it is unlikely one can purify Polδ4; and earlier reported purified and functionally characterized Polδ4 holoenzyme could very well be Polδ5 complex (11,15,23,24,39). Considering our both *in vivo* and *in vitro* analyses, we conclude that intrinsically human Polδ is a pentameric complex possessing two p12 subunits.

**Figure 6:**
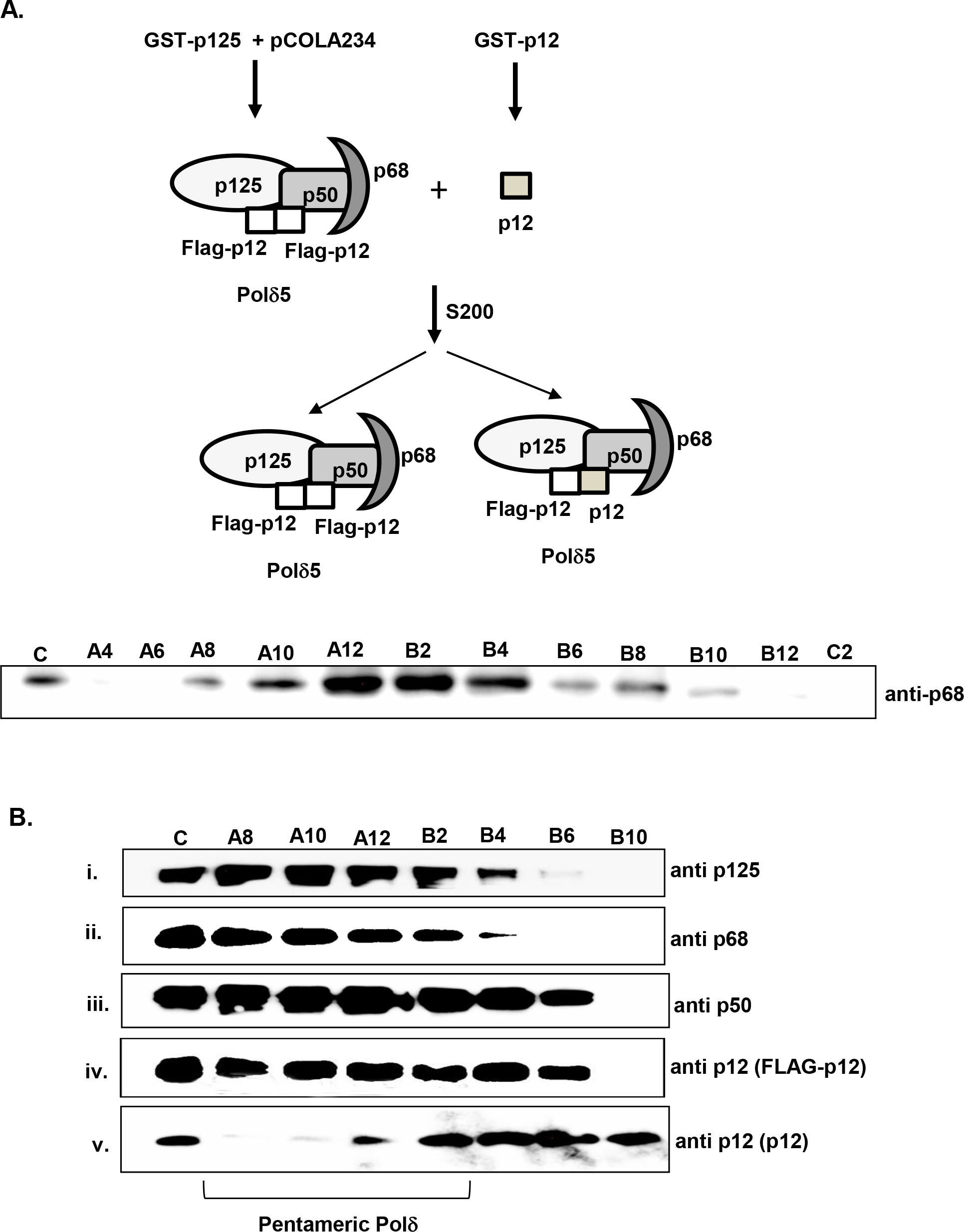
Purification of pentameric hPolδ holoenzyme. Schematic representation of purification of pentameric Polδ was shown. Mixture of Polδ4 and p12 was separated by gel filtration chromatography and various fractions was first analyzed by probing with anti-p68 antibody (**A**). Enriched fractions were again separated in SDS-PAGE, transferred to PVDF membrane. Membrane was cut into pieces as per the molecular weight of various subunits and were individually probed with specific antibody (**B**). Fractions possessing Polδ5 were denoted. Name of the fractions were as per the collections in 96 well plate.

### Dimerization of p12 is essential for PCNA interaction

Since, both R_3_K_4_R_5_ and _98_QCSLWHLY_105_ motifs are involved in PCNA interaction and the former motif additionally involved in protein dimerization; to ascertain any regulatory role of this motif in p12 function, a PCNA overlay experiment was carried out. Proteins were resolved on native PAGE to keep the natural folding and dimer structure intact as in Fig. 4A, transferred to PVDF membrane and blocking was performed in the presence of PCNA. After several washings, the blot was developed with anti-PCNA antibody (Fig. 7 A). As depicted in the figure, although p12 and its L104A, Y105A mutant formed dimers (upper panel, compare lane 1 and 3), signal for only wild type was detected (lower panel) suggesting that _98_QCSLWHLY_105_ motif is a true PIP motif required for PCNA interaction. Despite retaining PIP motif in the extreme c-terminal tail, due to its inability to form dimer, R3A, K4A, R5A mutant failed to bind to PCNA. Similarly, a pull down experiment was also carried out by taking a mixture of stoichiometric equivalents of purified GST-p12 or various p12 mutants and PCNA in Tris-buffer containing 150 mM NaCl salt concentration (Fig. 7 B). The mixture was incubated with GST-beads at 4 °C for 3 hrs in rocking condition, further beads were washed thrice and bound PCNA was eluted by SDS containing sample buffer. The eluted PCNA was detected by anti-PCNA antibody only when it was mixed with wild type p12 but not with L104A, Y105A or R3A, K4A, R5A mutant (compare lane 3 with 6 and 9).

**Figure 7:**
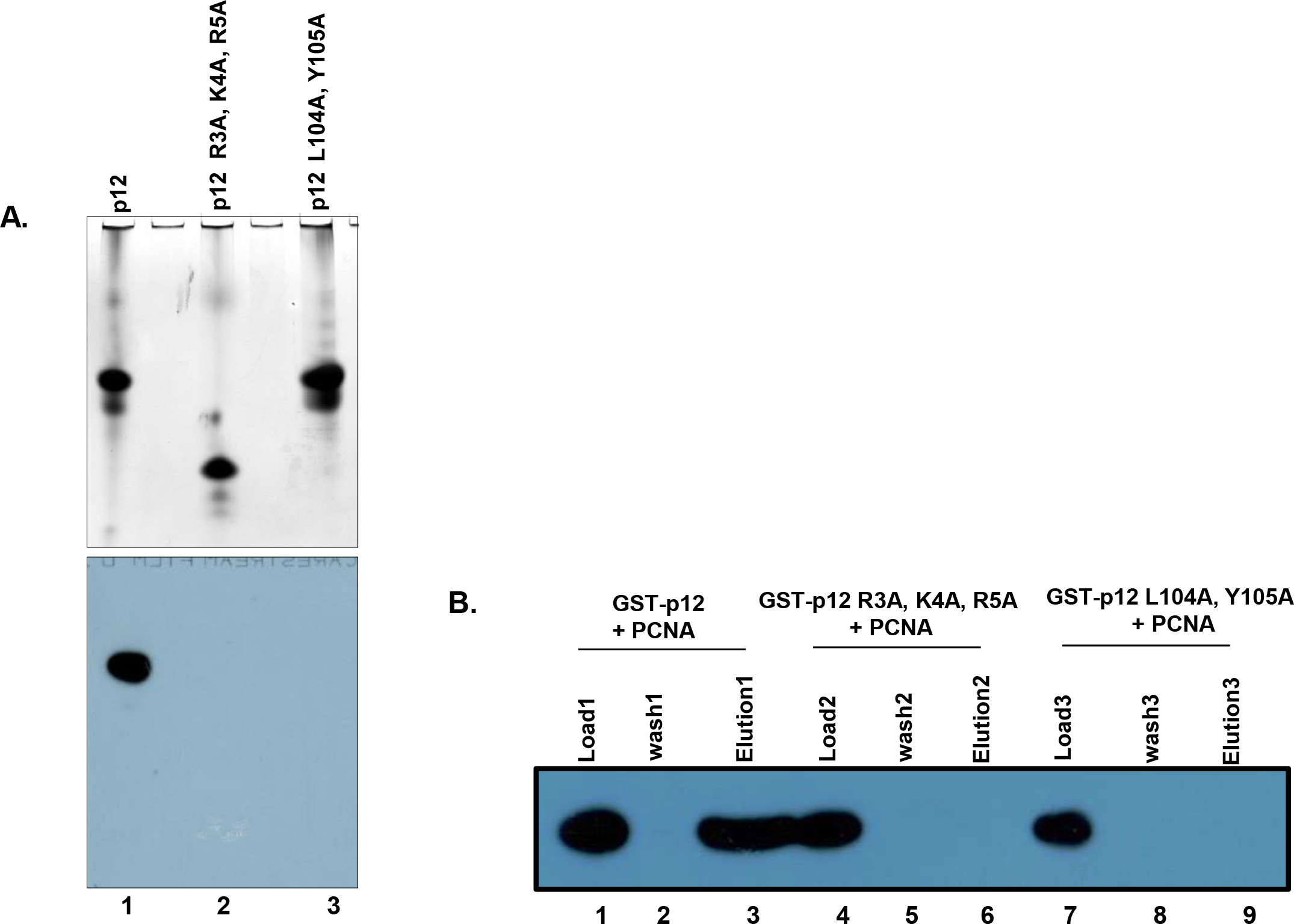
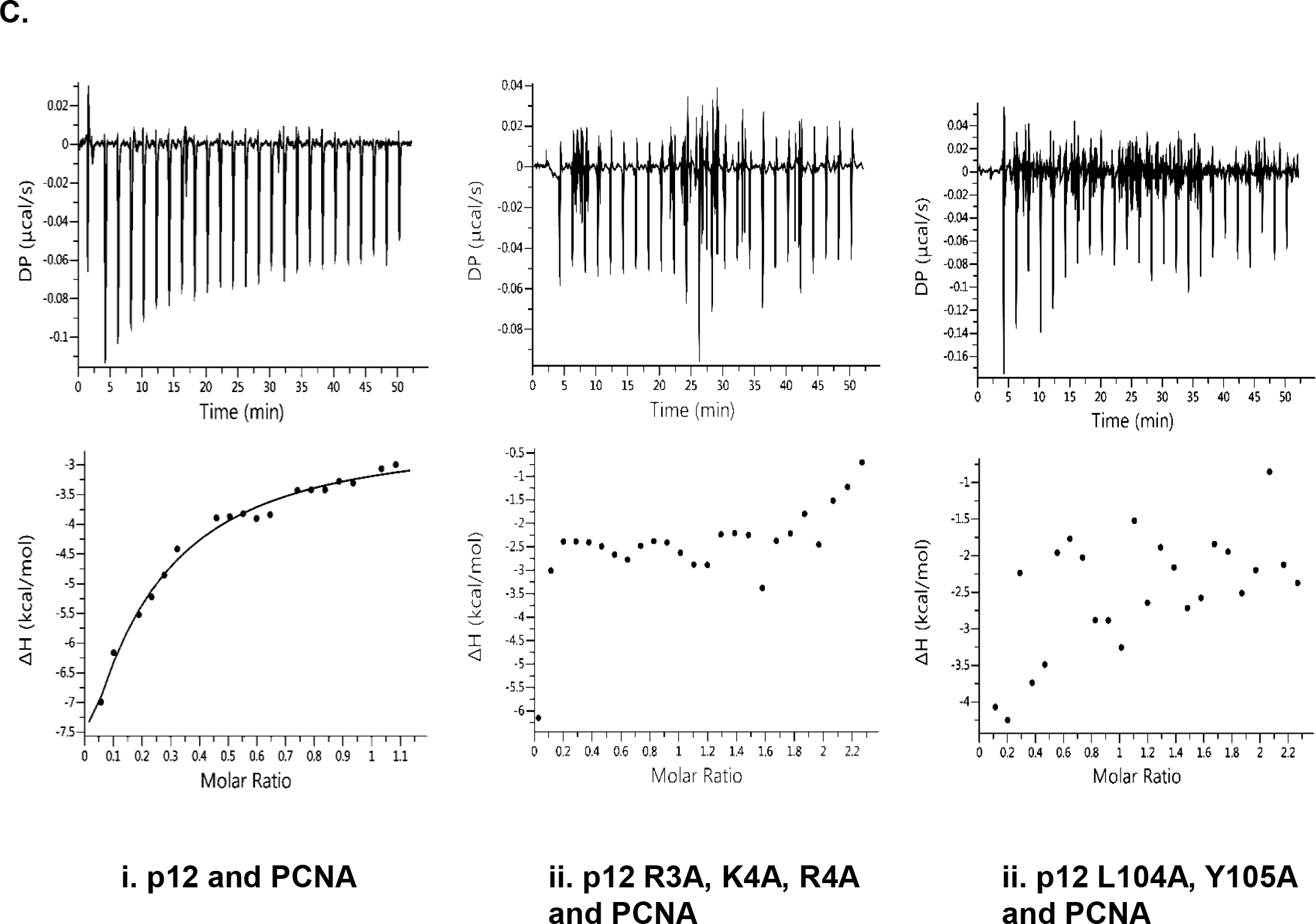
Mutations RKR motif inhibit binding with PCNA. **A.** Upper panel depicting coomassie blue stained gel of various p12 proteins resolved in a non-denaturating condition; whereas lower panel is a far-Western analysis of similar gel. Proteins were transferred from the gel to the membrane and further the blot was blocked with PCNA. After washings, bound PCNA was detected by anti-PCNA antibody. Lane 1, wild type; Lane 2, R3A, K4A, R5A; and Lane 3, L104A, Y105A p12 proteins. **B.** GST-pull down of PCNA by wild type p12. Beads of GST-p12 (lanes 1-3) or GST-R3A, K4A, R5A (lanes 4-6) or GST-L104A, Y105A p12 mutants (lanes 7-9) were mixed with PCNA in equilibration buffer, after incubation beads were washed and bound PCNA was eluted by protein loading dye. Various fractions were resolved in 12 % SDS PAGE, blotted to the membrane and developed by anti-PCNA antibody. Lanes 1, 4 and 7 are 10 % of Load; Lanes 2, 5 and 8 are 10 % of third washings; Lanes 3, 6 and 9 are the total elutes. **C.** ITC analysis of binding of wild type (i) or R3A, K4A, R5A (ii) or L104A, Y105A mutant (iii) p12 to PCNA at 25°C. In each panel, the upper half shows the measured heat exchanges during each PCNA protein injection. The lower half of each panel shows the enthalpic changes as a function of the molar ratio of the two proteins where p12 was considered as a dimer and PCNA as a trimer.

To determine the binding affinity of p12 with PCNA, ITC assay was carried out by placing p12 (10 μM) in the sample cell and PCNA (120 μM) was injected. The ΔH was ~ −80 kcal/mol, ΔG was ~ −6.80 kcal/mol, and the *K_D_* for the complex was 10 μM (Fig. 7 C). In similar assay condition, no significant heat exchange was observed while RKR- or PIP-motif p12 mutants was kept in the cell and PCNA in the syringe, suggesting no interaction between the proteins. Thus, our results suggest that dimerization at R_3_K_4_R_5_ motif promotes p12 interaction with PCNA via _98_QCSLWHLY_105_ motif.

### Inter-domain connecting loop region of hPCNA mediates its interaction with p12

In view of the fact that our predicted model structure showed p12 PIP peptide binding to IDCL domain of hPCNA; for confirmation, yeast two hybrid assay was carried out with p12 fused to Gal4 binding domain and two PCNA mutants namely, pcna-79 and pcna-90 fused to Gal4 activation domain. In pcna-79, two key hydrophobic residues L126 and I 128 of inter-domain connecting loop were mutated to alanines; whereas pcna-90 possesses two mutation P253,K254-AA in the extreme C-terminal tail of PCNA. Most of the interacting proteins bind to any of this two region of a trimeric PCNA ring (37). While wild type and pcna-90 were able to interact with p12 as evident from growth on SDA plate lacking leucine, uracil and histidine (Fig. 8 A, sectors 1 and 3); pcna-79 did not support the survival as it failed to form intact Gal4 by interacting with p12 (sector 2). p12 PIP mutant was used as a −ve control (sector 4). To strengthen our finding, GST pull down assay was carried out. Equal stochiometry of wild type and IDCL mutant of PCNA proteins were incubated with GST-p12, and pull-down assays were performed on glutathione-Sepharose affinity beads as described previously (39). In accordance with the data shown in Fig. 8 B, while GST-p12 was able to pull down most of the wild type PCNA from solution (compare lane 1 with 3), it failed to bind pcna-79 protein (compare lane 4 with 6) as detected by anti-PCNA antibody.

**Figure 8:**
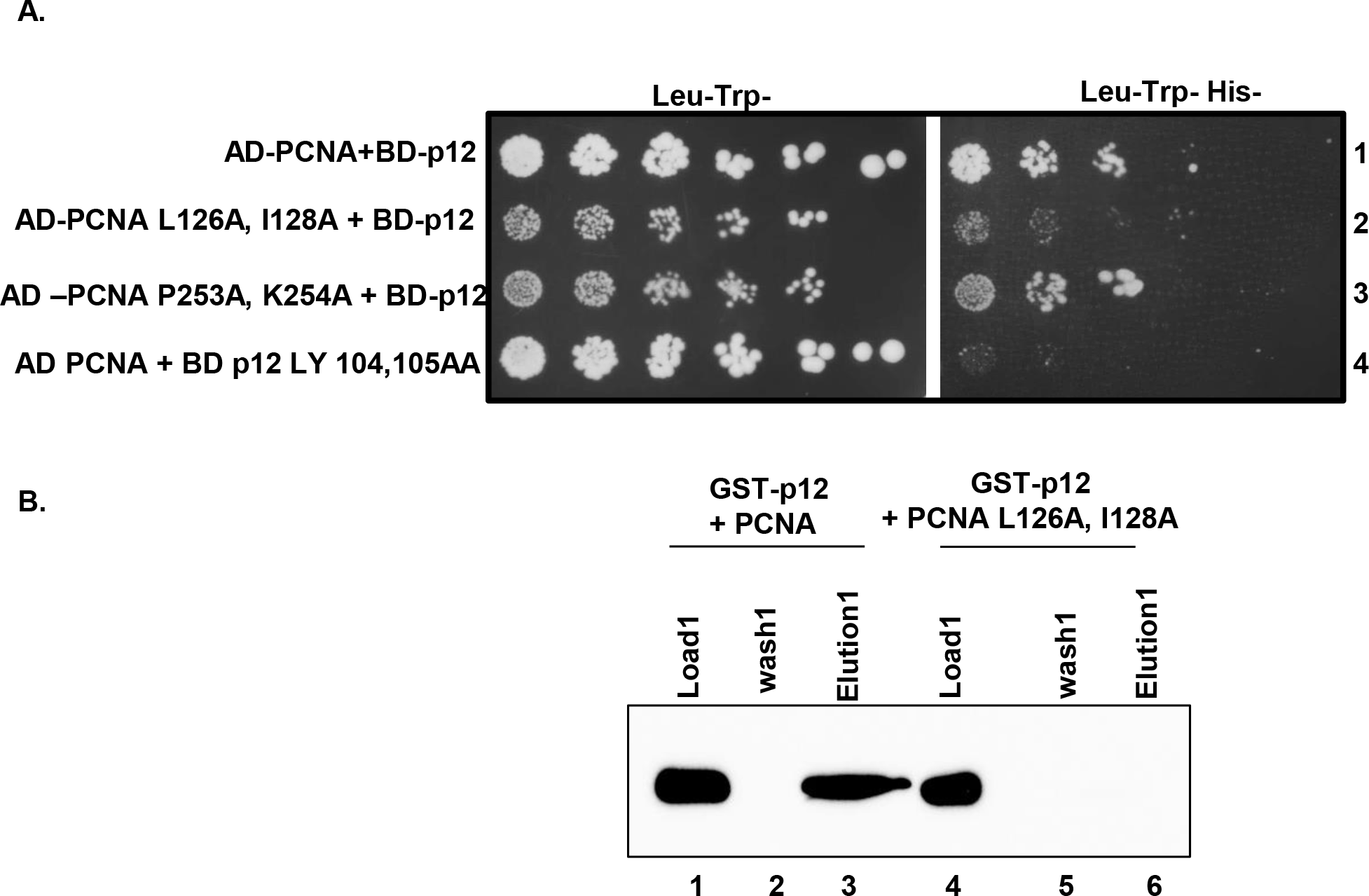
IDCL of hPCNA binding to p12. **A.** Yeast two hybrid analysis showing interaction of PCNA with p12. HFY7C yeast transformants with various GAL4-AD and BD fusions were selected on SD media plates lacking leucine, tryptophan with and without histidine amino acids. Row 1, AD-PCNA + BD-p12; Row 2, AD-PCNA L126A, I128A + BD-p12; Row 3, AD-PCNA P253A, L254A + BD-p12; Row 4, AD-PCNA + BD-p12 L104A, Y105A. **B.** GST-pull down of PCNA by wild type p12. Beads of GST-p12 were mixed with wild type (lanes 1-3) or L126A, I128A mutant PCNA (lanes 4-6) in equilibration buffer, after incubation beads were washed and bound PCNA was eluted by protein loading dye. Various fractions were resolved in 12 % SDS PAGE, blotted to the membrane and developed by anti-PCNA antibody. Lanes 1 and 4 are 10 % of Load; Lanes 2 and 5 are 10 % of third washings; Lanes 3 and 6 are the total elutes.

### Cdm1, a p12 homologue of *Schizosaccharomyces pombe* also forms dimer

The other fourth subunit of DNA polymerase delta that has been well characterized is Cdm1, a p12 orthologue from *S. pombe* (10). Cdm1 consists of 160 amino acids, MW of 18.6 kDa and pI of 7.73. Like p12, it shows abnormal mobility in the SDS PAGE and migrates as ~22 kDa protein (Fig. 9 A). As it also has conserved RKR-motif (K_2_K_3_R_4_ in Cdm1) at its N-terminal end, we wanted to examine whether homodimerization property of the smallest subunits of Polδ is evolutionarily conserved. Wild type and K2A, K3A, R4A mutant Cdm1 proteins were purified to near homogeneity from bacterial over-expression systems and analyzed on native PAGE. Just like p12, the wild type Cdm1 migrated slower than its mutant. The slower migration of Cdm1 complex comparison to p12 dimer could be attributed to the differences in their pI and MW (pI 7.3 *vs* 6.3; MW 12 *vs* 18.6). The dimeric p12 incidentally migrates at about the same position with monomeric Cdm1 mutant protein (Fig. 9 B). Thus, we concluded that homodimerization is the intrinsic property of the 4^th^ subunits of Polδ and is mediated by a stretches of conserved basic amino acids located at the extreme amino terminal end (RKR/KKR-motif).

**Figure 9:**
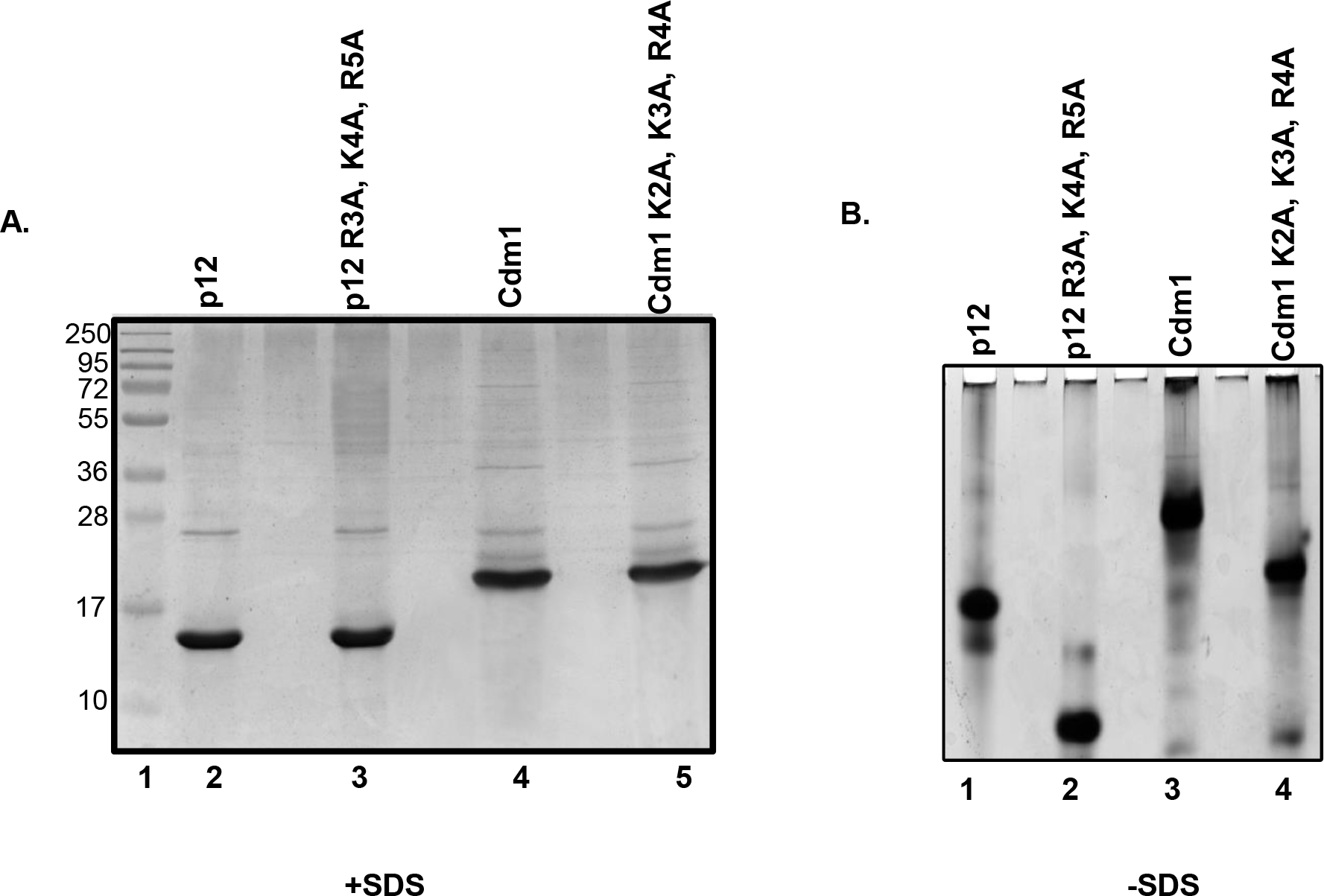
Cdm1 also dimerzes via KKR-motif and stability of the dimers. **A.** Purity of wild type and K2A, K3A, R4A mutant Cdm1 proteins were analyzed in 12 % SDS-PAGE and their mobility was compared with p12 proteins. **B.** These proteins were further resolved in PAGE without SDS. Lane 1, p12; Lane 2, p12 with R3A, K4A, R5A mutations; Lane 3, Cdm1; and Lane 4, Cdm1 with K2A, K3A, R4A mutations.

## Discussion

DNA polymerase δ is a high fidelity essential DNA polymerase not only plays a central role in DNA replication, it also participates in DNA recombination and several DNA repair pathways from yeasts to human (40,41). Several mutations in the mouse and human Polδ subunits have been mapped to cause various cancers and genome instability in yeasts (42–44). Thus, it is important to understand function of each subunits and their precise role in processivity and fidelity of the holoenzyme. In this report, we have re-investigated role of the smallest subunit of hPolδ, p12 in holoenzyme architecture and PCNA interaction.

As reported earlier and also in this study p12 subunits interact with p125 and p50, whereas p50 makes a connecting bridge between catalytic p125 and accessory p68 subunits (45). Thus, hPolδ is proposed to be a heterotetrameric holoenzyme. Our yeast two hybrid assay, cellular co-localization in replication foci, mutational analyses and several physio-biochemical assays including formaldehyde crosslinking clearly demonstrate that p12 exists as a homodimer both *in vivo* and *in vitro;* and dimerization is dependent on amino terminal tripeptide basic amino acids sequence, _3_RKR_5_-motif. It also argues against the tetrameric nature hPolδ. As it has been previously shown, various sub-assemblies of hPolδ holoenzymes could exist in the cell based on Polδ’s function in either replication or repair (46), we propose that among these sub-complexes pentameric Polδ is the native form of Polδ. Pull-down of cellular Polδ by a tagged p12 and *in vitro* reconstitution of Polδ5 definitely authenticates our prediction (Fig. 2 A and Fig. 6). Purification or *in vitro* reconstitution of Polδ holoenzymes by several groups also indicated higher stochiometry of p12 in comparison to other subunits in Polδ (19,25,38,45–47). We also found that the dimerization of the 4^th^ subunit of Polδ is not restricted to human as Cdm1 of SpPolδ also forms a dimer that is again dependent on KKR motif. As the RKR/KKR motif has been retained in other p12 homologues as well, it appears that such a property of the smallest subunit of Polδ is evolutionarily conserved. Interestingly, the small accessory subunit of yet another B-family polymerase Polζ (a complex of Rev3 and Rev7) Rev7 is found to function as a dimer (48). Thus, subunit dimerization of B-family DNA pols could be an intrinsic property; and it will be advantageous for the DNA polymerases to establish multiligand interactions during replication.

Most of the PCNA interacting partners including Polδ bind to PCNA through a structurally conserved canonical PIP sequences (49). The PIP sequences structurally organized into a 310 helix that functions as a “hydrophobic plug” which fits into the IDCL domain of PCNA. The affinity of PIP sequences depend on how snuggly the 3_10_ helixes fit into the binding surface of PCNA. Previously, we have shown that all the three subunits of ScPolδ contribute to PCNA interaction as well as its processive DNA synthesis (5). The PIP motifs in Pol3, Pol31 and Pol32 have been mapped and each of them can bind to each monomer of trimeric PCNA. The complication arises in hPolδ complex and it’s not clear which subunits primarily contribute to PCNA interaction and processivity of the enzyme. Although, reports have shown that all the four subunits bind to PCNA and contribute enzymatic processivity to different degrees, identification of PIPs remains elusive (15). PIP motif in the largest subunit of hPolδ, p125 is yet to be deciphered. Like Pol32 subunit, p68 possesses a conventional PIP motif at the extreme carboxyl terminal end (49). Therefore, deletion of last 20 amino acids of p68 that encompass PIP sequences failed to bind to PCNA. Another study also revealed a mechanism that could modulate the interaction of p68 with PCNA by a protein kinase-mediated phosphorylation of Ser458 in the PIP-box _456_QVSITGFF_463_. Accordingly, it was proposed that phosphorylation/dephosphorylation of the PIP motif of p68 could provide a switch for the reorganization of replication proteins on the PCNA−Polδ complex (50). Although the PIP-box is commonly found near the C-terminus, some proteins such as Pol31, RFC, DNA-cytosine-5-methyltransferase, and WRN DNA helicase, have internal or N-terminal PIP-boxes. Similarly, a PCNA interaction domain for p50 was identified at the N-terminus. A 22-amino acid oligopeptide containing the PIP sequence _57_LIQMRPFL_64_ was shown to bind PCNA by far Western analysis (19,20); however, a mutational analysis in this motif to provide functional evidence is yet to be carried out. Crystal structure of p50 reveals that _57_LIQMRPFL_64_ sequences belong to α2 helix of the *OB*-fold generally involved in DNA binding and is quite different from the commonly found 3_10_ helix structures in PIP motifs (51). Extensive biochemical studies have been carried out earlier to suggest that absence of p12 impedes processive DNA synthesis of Polδ. Here, by using structural modeling, yeast two-hybrid system and many other biophysical and chemical studies we have mapped PIP motif of p12 at the C-terminal tail that forms a typical 3_10_ helix that interacts with IDCL of PCNA. Additionally, we showed that while C-terminal motif _98_QCSLWHLY_105_ is a PIP motif that is directly involved in PCNA interaction, the N-terminal motif _4_KRLITDSY_11_ is involved in dimerization and participates in regulating p12 interaction with PCNA. Contrary to earlier observation, our study reveals that the role of RKR-motif in PCNA interaction is mostly indirect (14). Thus, like phosphorylation/dephosphorylation of p68, dimerization could be yet another mode of regulation by p12 subunit to control switch in interacting partners by Polδ with PCNA. In addition to dimerization, change in p12 protein turnover during genotoxic stress is yet another mode of Polδ functional regulation (52,53).

There are sufficient evidences available both *in vitro* and *in vivo* to support an idea that multiple subassemblies of Polδ may exist. Proteolysis of p12 and p68 subunits by human calpain-1 could trigger interconversion of Polδ in the cell. Considering dimerization of p12 and purification of Polδ5, we propose existence of four different complexes of Polδ; Polδ5 (p125+p50+p68+2xp12), Polδ3 (p125+p50+p68), Polδ2 (p125+p50) and p125 alone (Fig. 8 A). Depending upon the cellular contexts such as cells either actively replicating or under genomic stress, this interconversion between the Polδs might be transpired. Polδ5 could be the major holoenzyme that takes part in DNA replication, which was earlier extensively studied as Polδ4. Polδ4 with a monomeric p12 does not exist in the cell. So, now instead of four subunits, five subunits of Polδ should be considered that will interact with three available IDCL in the trimeric PCNA unless they use other binding sites such as inter-subunit junction or C-terminal domain of PCNA (54,55). Genetic analyses of Polδ PIPs in *S. cerevisiae* revealed that for cell survival, along with Pol32 PIP, any one Pol3 or Pol31 PIP are essential. In the absence of functional Pol32 PIP domain, PIP domain mutation in Pol3 or Pol31 subunits causes lethality(5). Despite being structural subunits, *CDC27* and p68, the Pol32 homologues are deciphered to be essential in *S. pombe* and mice, respectively (56); it explains their major role in Polδ’s function in DNA synthesis. Binding affinity of Polδ with PCNA also increases when p12 or p68 binds to the core (*Kd*= 8.7~9.3nM), and it further increases when all the subunits are present (*Kd*= 7.1nM). The *Kd* of Polδ core is found to be 73 nM (45). Accordingly, the addition of p68/Pol32 to core (p125+p50 or Pol3+Pol31) results in high processivity, thus its binding to PCNA appears to be critical. Considering all these, we propose a model for the network of protein-protein interactions of the Polδ-PCNA complex (Fig. 10). In a pentameric state of hPolδ, along with p68, any other two subunits among p125/p50/p12 will bind to PCNA in any combinations as shown in the figure 10 B (i to iv). Upon p68 degradation or its phosphorylation, p125, p50 and one monomer of p12 dimer can bind to PCNA (v). Similarly, upon p12 proteosomal degradation as a response to DNA damage; other three subunits will make contacts with PCNA (vi); whereas due to cleavage of both p68 and p12 in certain situation, p125 and p50 bind to PCNA (vii), although the Polδ core functions with low processivity. Thus, this study warrants extensive mutational analyses as had been carried out in yeast Polδ holoenzymes rather than analyzing sub-complexes to decipher precise role of these PIPs in cellular function and processive DNA synthesis by hPolδ.

**Figure 10:**
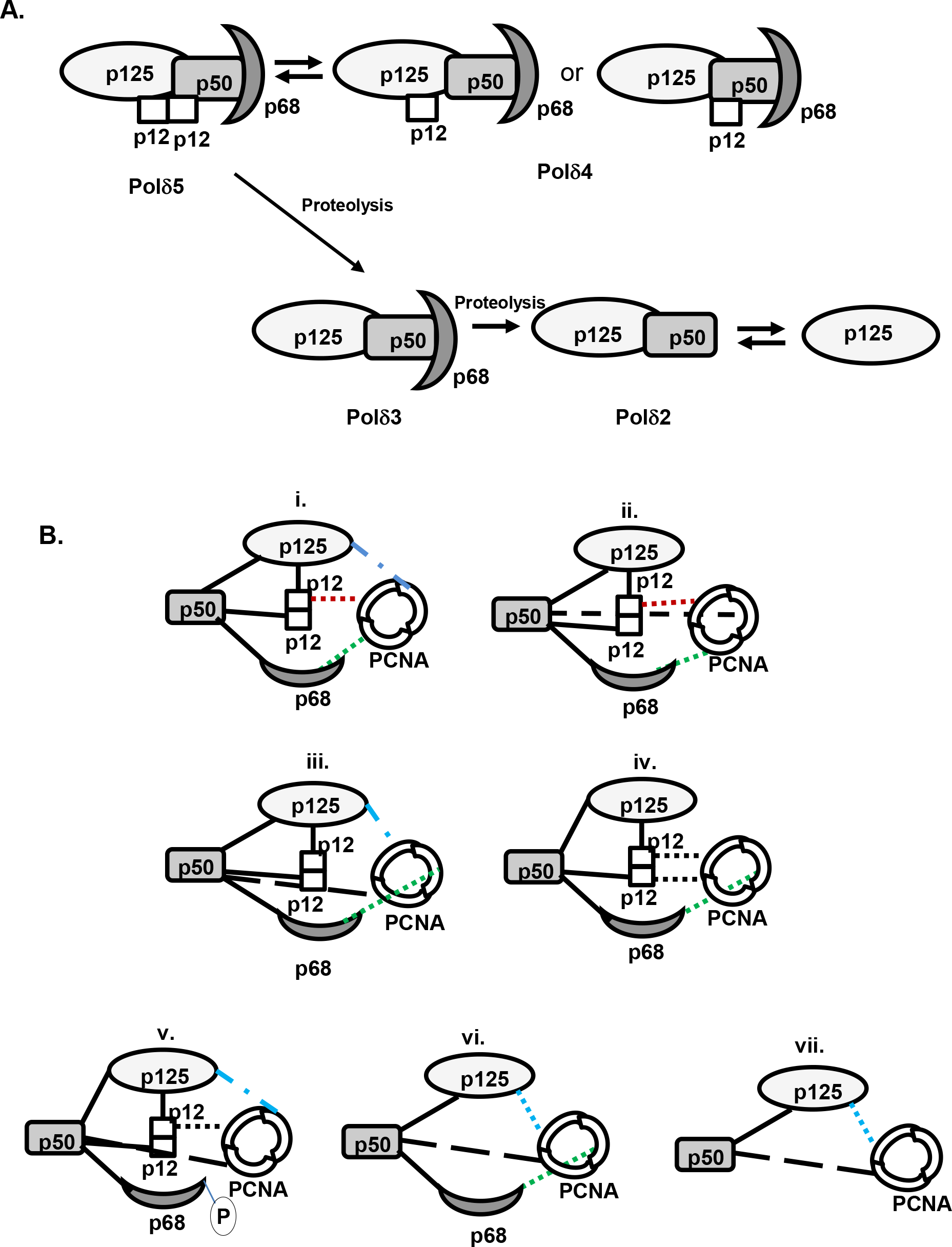
Protein–protein network model for hPolδ and PCNA. **A.** Depicting different sub-assemblies of Polδ complexes. As p12 is a dimer, pentameric Polδ holoenzyme is proposed to function in DNA replication. Polδ4 unlikely exists in the cell; however, after proteolysis during certain conditions like genomic stress; Polδ5 can be downgraded to Polδ3 or Polδ2 complexes. **B.** Four different proposed modes of Polδ5 binding to PCNA where apart from p68, any other two subunits can bind to IDCLs of PCNA. In dimerization state only p12 can bind to PCNA (i–iv). Upon phosphorylation of p68 or proteolysis of p12, other remained three subunits bind to PCNA (v and vi). In case of core, both subunits interact with PCNA but with compromised processive DNA synthesis. PCNA monomer binding to p125, p50, p68 and p12 are shown in blue, black, red and green dotted lines, respectively.

In conclusion, here we show that RKR-mediated dimerization plays a vital role in p12 binding to PCNA and Polδ5 architecture, and the phenomenon appears to be conserved throughout evolution. Dimerization of p12 could play an instrumental role in re-orchestrating pentameric Polδ-PCNA complex at the replication fork and thereby it regulates Polδ’s function.

## Experimental Procedures

### Plasmids, oligonucleotides, antibodies and enzymes

Human DNA polymerase delta constructs pCOLA-hPold234 and pET32-p125 (kind gift from Prof. Y. Matsumoto) were used as precursor plasmids for subsequent manipulation (57). The oligonucleotides (Supp. Table-1) from Integrated DNA Technologies (IDT, USA), Q5 high fidelity DNA polymerase and other restriction enzymes from NEB, and antibodies from Sigma or Abcam were procured. Human PCNA was directly amplified from c-DNA synthesized from total RNA of HELA cell line by using primers NAP239 and NAP240, and cloned into pUC19 vector. PCR product was digested with BamHI and cloned into BglII site to generate a bacteria expression system of GST-fused hPCNA. To express in yeast, hPCNA was amplified by using primers NAP251 and NAP240, PCR product was digested with BamHI and cloned into same site of pGBT9 and pGAD424 to generate Gal4BD-hPCNA and Gal4AD-hPCNA, respectively. Inverse PCR was carried out using primer set NAP300-NAP304 on pUC19-hPCNA to generate pcna-79 (L126,128AA); and further subcloned into any expression vector. Human pcna-90 (P253A, K254A) was PCR amplified using primers NAP251 and NAP305 from pUC19-hPCNA template, PCR product was digested BamHI and cloned into same site of pUC19 and pGBT9. A GFP-fused hPCNA expression construct under CMV promoter was a gift from Prof. Wim Vermeulen.

Amplified PCR products of wild type, R3A, K4A, R5A; and L104A, Y105A mutants of p12 from pCOLA-hPold234 by using primer pairs NAP261-NAP262, NAP362-NAP261, and NAP265-NAP262, respectively; were digested with EcoRI-BamHI and cloned into same sites of pGBT9 and pGAD424. Similar PCR products amplified using primers NAP260-NAP261, NAP373-NAP261, and NAP265-NAP260 were also digested alone with BamHI and cloned into BglII site of either pNA716 or p3X Flag CMV7.1 vector to generate N-terminal GST tag proteins expressed under T7 promoter and N-terminal FLAG-tag human cell expression system, respectively. For Confocal microscopy and pull down studies, BamHI fragments of various p12 were subcloned into BamHI site of pcDNA-GFP vector. For RFP-fusion, p12 fragment and catalytic domain of Polθ were amplified with the primer sets NAP448-NAP450 and NAP444-NAP151, respectively; digested with EcoRI-BamHI and cloned into same sites of pASred2-c1 vector.

Other hPolδ subunits such as p125, p50 and p68 were also PCR amplified by the primer sets NAP252-NAP248, NAP254-NAP255, and NAP258-NAP257, respectively; digested with EcoRI-BamHI and cloned into same sites of pGBT9. Wild type and K2A, K3A, R4Amutant of Cdm1 were PCR amplified from *S. pombe* genomic DNA using primers NAP361-NAP451 and NAP361-NAP452, respectively; and the BamHI digested products were cloned into BglII site in pNA716 for bacterial expression. Similarly, GST-hp125 expression plasmid was generated by cloning a BamHI digested PCR product amplified from pET32p125 as a template using NAP247 and NAP248 primers into BglII site of pNA716. All of these constructs were authenticated by DNA sequencing.

### Protein Purifications

All GST tagged proteins were expressed in *Escherichia coli BL21 DE3* and purified by affinity chromatography using glutathione sepharose beads (GE-healthcare). The proteins were expressed as amino terminal GST-fusion proteins under T7 promoter. Briefly, 5 ml pre-culture of the transformant was added to 500 ml LB + 50 μg/ml ampicillin and grown at 37 °C till the OD_600_ reaches to 0.6. Next the culture was induced with 1 mM IPTG and allowed to grow for another 8 hrs. Cells were harvested and about 3 gm of frozen cells were resuspended in 1X cell breaking buffer (50 mM Tris–HCl pH 7.5, 10 % sucrose, 1 mM EDTA, 500 mM NaCl, 0.5 mM PMSF, 0.5 mM Benzamidine hydrochloride, 10 mMβ-mercaptoethanol and protease inhibitor cocktail). Cells were lysed with a high pressure homogenizer at 10K psi (STANSTED). The lysate was cleared after centrifugation at 10K rpm for 10 min. Further the supernatant was centrifuged at 30K rpm for 1 hour in P70AT rotor (Hitachi). Rest of the steps used for purification was same as described before (39). All the proteins were stored in the buffer containing final concentration of 50 m*M* Tris–HCl, pH 7.5, 150 m*M* NaCl, 10% glycerol, 5 m*M* dithiothreitol (DTT) and 0.01% NP-40. The purity of the protein was confirmed after resolving on 12% SDS-PAGE and stained by Coomassie Blue.

However, for human Polδ purification, bacterial strain BLR (DE3) was co-transformed with pLacRARE2 plasmid from Rosetta2 strain (Novagen), GST-p125 and pCOLA-hPold234; and colonies were selected on LB agar plate containing ampicillin (50μg/ml), kanamycin (30μg/ml) and chloramphenicol (35μg/ml). About 60 ml overnight grown pre-culture was inoculated into 6 L LB with mentioned antibiotics and grown at 37 °C to an OD600 of 0.6;followed by induction with 1 mM IPTG and further growth was continued for 15 hrs at 16 °C. The cells were harvested and stored at −80°C until use. The cell breaking condition and other purification steps were followed as mentioned. Taking advantage of strategically located PreScision protease site, cleaved Polδ4 (p125-p50-p68-p12) was obtained in which only p12 subunit was amino-terminally FLAG tagged. Similarly, p12 protein was also purified by using bacterial GST-p12 construct.

### Size exclusion Chromatography

For size exclusion chromatography, about 10μg of each purified p12 proteins were loaded onto a Superdex 200 PC3.2/30 minicolumn pre-equilibrated with buffer containing 50mM HEPES pH7.5, 150 mM NaCl and 10% glycerol. Chromatography was performed on an AKTApure M system (GE Healthcare) at a flow rate of 0.05 ml/min at 4°C, and the absorbance was monitored at 280 nm.

### Purification of human Polδ5 complex

In order to purify Polδ5 complex; purified Polδ4 (10μg) was mixed with equal amount of untagged p12 protein under rocking conditions for 4 hrs at 4°C. Then the mixture was injected in Superdex 200 PC3.2/30 minicolumn pre-equilibrated with buffer containing 50mM HEPES pH7.5, 150 mM NaCl and 10% glycerol. Chromatography was performed on an AKTApure M system (GE Healthcare) at a flow rate of 0.03 ml/min at 4°C, and the absorbance was monitored at 280 nm. The elute was collected in a 96 well plate fraction collector and then the various fractions were subjected to SDS-PAGE and western blot for the detection of individual subunits of DNA polymerase delta. To check the presence of Polδ fractionation, membrane was first probed with anti-p68 antibody (Cat # WH0010714M1) since it does not directly interact with p12 and will give a clear indication of the complex. Further, the enriched fractions were analyzed by probing with specific antibody such as anti-p125 (Cat # SAB4200053), anti-p50 (Cat# SAB4200054), and anti-p12 (Cat # WH0057804M1). Horseradish peroxidase-conjugated host specific secondary IgG (Cat# A90376154; Sigma–Aldrich and Cat# W402B, Abcam) was used to develop the blot by Chemidoc.

### Yeast two hybrid analyses

The yeast two-hybrid analyses were performed using *HIS3* as a reporter system(39). The HFY7C yeast strain (from Clonetech) was transformed with various combinations of the *GAL4*-AD (*TRP1*) and -BD (*LEU2*) fusion constructs. Co-Transformants were obtained on synthetic dropout (SD) media plates lacking leucine and tryptophan. In order to verify the interaction, transformants were grown on 5 ml YPD liquid medium over night at 30° and various dilutions were either streaked or spotted on Leu^−^Trp^−^His^−^ selection medium. Further the plates were incubated for 2 days at 30 °C and photographed. Yeast transformants exhibiting histidine prototrophy are indicative of protein-protein interaction.

### Formaldehyde cross-linking

About 1-5 μg of native or mutant p12 in 20 mM HEPES buffer (pH 7.5) was mixed with 0.5% formaldehyde solution for 30 min at 25°C. The reaction was terminated by the addition of SDS sample buffer. Cross-linked proteins were resolved by electrophoresis in a 12 % SDS-PAGE. The gel was stained with coomassie blue.

### Native PAGE analysis

Wild type and various mutants of p12 and Cdm1 proteins were mixed with DNA loading dye and were analyzed on 12 % Native PAGE. The gel was run at 80 volts for 5-15 hrs at 4°C using running buffer (25 mM Tris-base, 192 mM glycine, pH 8.8). The proteins were visualized by Coomassie Blue staining of the gels.

### Confocal Microscopy

CHO cells were grown up to 70 % confluency on cover glass in DMEM media supplemented with 10% FBS and 1X penicillin-streptomycin antibiotics solution. Further the cells were washed with DPBS, pH 7.4 and then replaced with DMEM media containing 5 % FBS. These cells were co-transfected with GFP/RFP fusion constructs of p12, PCNA and a catalytic domain of Polθ in various combinations as required by using Lipofactamine transfection kit as per the manufactures’ protocol (Invitrogen). Further, cells were incubated at 37 °C with 5 % CO_2_ and 95 % relative humidity. After 48 hrs, cell were thoroughly washed thrice with DPBS, fixed with ice cold 100 % methanol at −20 °C for 20 mins; followed by rinsing with DPBS, pH7.4, and then slide were prepared using antifade as mounting agent. Images were taken using Leica TCS SP5 at 63 X objective.

### Co-immunoprecipitation

HEK293 cells were grown up to 70% confluency in 10 cm dish containing DMEM media supplemented with 10% FBS and 1 X penicillin – streptomycin antibiotics. These cells were co-transfected with FLAG-p12 with either GFP-p12 or GFP-p12 R3A, K4A, R5A mutant by using Lipofactamine transfection kit. Cells were grown in humidified CO2 incubator at 37°C. After 48 hrs growth, cell were harvested, washed thrice with DPBS and immediately resuspended in RIPA buffer (50 mM Tris-HCl pH 8.0, 0.5 % Sodium deoxycholate, 1000 mM NaCl, 0.1% SDS, 1mM EDTA, 1mM EGTA, 25 mM Sodium Pyrophosphate, 1 mM β-glycerophosphate, 1 mM sodium orthovandate and protease inhibitor tablet) and kept for 1 hour at 4 °C rocking condition. Followed by centrifugation at 10000 rpm, the supernatant was collected and protein concentration was determined using Bradford method. About 500 μg of total protein was incubated overnight with anti-FLAG antibody conjugated agarose beads. The beads were washed thrice with RIPA buffer and bound proteins were eluted by 40 μl of SDS loading buffer and subjected to 12 % SDS PAGE. The proteins from the gel were transferred to PVDF membrane, followed by incubation of the membrane with 5 % skim milk in PBST for 1 hour at room temperature. The blot was washed thrice with PBST and incubated with anti-GFP antibody (1:5000 dilution, cat# ab290 from abcam) for 2 hours at RT. Subsequently, after through washings, horseradish peroxidase-conjugated goat anti-rabbit IgG (diluted 1:10000 in PBST, cat# A6154; Sigma-Aldrich) was used to develop the blot.

Similarly, native human Polδ was co-immuno-precipitated from HEK293 cells transfected with GFP-p12. About 500 μg of total protein was incubated overnight with anti-GFP or anti-p125 antibody conjugated agarose beads. The beads were washed thrice with RIPA buffer and bound proteins were eluted by 40 μl of SDS loading buffer and subjected to 12 % SDS PAGE. The proteins from the gel were transferred to PVDF membrane, cut into 4 pieces as per the molecular weight markers, membranes were individually incubated with 5 % skim milk in PBST for 1 hour at room temperature. The blot was washed thrice with PBST and probed with subunit specific antibody for 2 hours at RT. For p50 probing, first the membrane was probed with anti-p50 antibody; then stripped off, and again probed with ant-GFP antibody as both the proteins migrate close to each other. Subsequently, after through washings, horseradish peroxidase-conjugated goat anti-IgG (diluted 1:10000 in PBST, cat# A6154; Sigma-Aldrich) was used to develop the blot.

### PCNA overlay Assay

Various proteins were resolved in two 12 % Native PAGE and while one of the gel developed with Coomassie blue, other one was transferred to methanol activated PVDF membrane. The blot was first washed with BLOTTO (25 mM Tris-HCl, pH 7.4, 150 mM NaCl, 5 mMKCl, 5 % fat-free milk, 1 % BSA, 0.05 % Tween 20) for 1 h at room temperature. Then, the blot was incubated overnight at 4 °C in 10 ug/ml of PCNA containing BLOTTO with constant agitation. After three rinses with BLOTTO, the membrane was incubated with anti-PCNA antibody (diluted 1:1000, cat#SAB2108448; Sigma-Aldrich) in BLOTTO. Subsequently, after through washings, horseradish peroxidase-conjugated goat anti-rabbit IgG (diluted 1:10000 in PBST, cat# A6154; Sigma-Aldrich) was used to develop the blot.

### GST pull down assay

GST-wild type or LY 104,105 AA p12 protein bound glutathione sepharose beads was mixed with 0.5 μg of either wild type or mutant (L126A, I128A) human PCNA and pull down experiment was carried out using a standardized protocol described previously (39). Then the beads were thoroughly washed three times with 10 volumes of equilibration buffer (50 mM Tris-HCl, pH 7.5, 150 mM NaCl, 5 mM dithiothreitol, 0.01 % NP-40, 10 % glycerol). Finally, the bound proteins were eluted with 50 μl SDS loading buffer. Various fractions were resolved on a 12 % SDS-PAGE; this was followed by Western blot, similarly performed as in co-immunoprecipitation except that the primary antibody used is anti-PCNA antibody (cat#SAB2108448; Sigma-Aldrich) in 1:750 dilutions.

### Isothermal titration calorimetry

The purified p12 and PCNA protein were dialyzed overnight in 1 liter of 150 mM sodium chloride and 20mM HEPES at pH 7.4 at 4°C. Final concentration of p12 was adjusted as 10 μM and PCNA as 120 μm. ITC were performed at 25°C using a Malvern MicroCal ITC calorimeter. The p12 was in the sample cell and the PCNA was in the syringe. Twenty five injections of the PCNA were done at intervals of 120 seconds and initial delay of 120 second. The data were analyzed to determine using a single-site binding model provided in the ITC analysis software package. Similarly, p12 oligomerization was checked by ITC at 30° C with twenty injections of protein.

### Circular Dichorism

The purified p12 protein were dialyzed overnight in 1 liter of 20 mM sodium chloride and 20 mM HEPES at pH 7.4 at 4°C.The secondary structure of p12 and p12 R3A,K4A,R5A mutant were determined by circular dichroism (CD) spectroscopy using a ChiraScan applied photophysics. Spectra were taken at 25 °C in a 10 mm path length quartz cuvette containing the sample at concentrations of 0.2 mg/ml of protein in 20 mM HEPES buffer, pH 7.5, 20mM NaCl. The spectra were corrected for buffer. Mean residue ellipticity values were calculated using the expression [θ] = θ×100 / (cln), where θ is the ellipticity (in millidegrees), c the protein concentration (in mol/liter), l is the path length (in cm), and n is the number of amino acid residues.

### *In silico* analysis of p12 structures

p12 RKR (1-MGRKRLITDSYPVK-14) and PIP (92-GDPRFQCSLWHLYPL-106) domains were used for peptide structure prediction by using PEP-FOLD3 server (http://bioserv.rpbs.univ-paris-diderot.fr/services/PEP-FOLD3/). The models generated were validated by SAVES and Ramachandran plot showed all residues in allowed regions, validating both the models. Further, the generated structural models were aligned with PIP peptide sequences from p21 (1AXC) and p68 PIP (1U76).

## Acknowledgements

We are grateful to Profs. Y. Matsumoto and W. Vermeulen for providing us human Polδ expression and CMV-GFP-hPCNA plasmids, respectively. We thank Sitendra Prasad Panda for his technical assistance, Dr. Jawed Alam for his involvement in initial studies, and Bhabani Shankar Sahoo for his help in confocal microscopy. Our laboratory colleagues are acknowledged for their thoughtful discussion. PK is a DBT-Senior Research Fellow; KM and DP are thankful to CSIR-SRF and DBT-RA fellowships, respectively. This work was supported by the intramural core grant from ILS, Bhubaneswar, India.

## Conflict of interest

The authors have no conflicts of interest.

## Author contributions

NA and PK designed the study; PK, DP and KM generated materials; PK carried out experiments; NA, PK, DP and KM analyzed the data and prepared the manuscript; NA conceptualized the study and acquired funding for the project. All the authors reviewed the results and approved the final version of the manuscript.

**Abbreviations**
Pol: DNA polymerase
PCNA: proliferating nuclear antigen
PIP: PCNA interacting peptide
SDS: Sodium dodecyl sulfate
PAGE: polyacrylamide gel electrophoresis
UV: ultra violet
HU: hydroxyurea
GFP: green fluorescence protein
RFP: red fluorescence protein
ER: endoplasmic reticulum
NLS: nucleus localizing signal

## References

1. Pavlov, Y. I., Shcherbakova, P. V., and Rogozin, I. B. (2006) Roles of DNA polymerases in replication, repair, and recombination in eukaryotes. International review of cytology 255, 41–132

2. Stillman, B. (2008) DNA polymerases at the replication fork in eukaryotes. Molecular cell 30, 259–260

3. Kunkel, T. A., and Burgers, P. M. (2014) Delivering nonidentical twins. Nature structural & molecular biology 21, 649–651

4. Johnson, R. E., Klassen, R., Prakash, L., and Prakash, S. (2015) A Major Role of DNA Polymerase delta in Replication of Both the Leading and Lagging DNA Strands. Molecular cell 59, 163–175

5. Acharya, N., Klassen, R., Johnson, R. E., Prakash, L., and Prakash, S. (2011) PCNA binding domains in all three subunits of yeast DNA polymerase delta modulate its function in DNA replication. Proceedings of the National Academy of Sciences of the United States of America 108, 17927–17932

6. Waga, S., Bauer, G., and Stillman, B. (1994) Reconstitution of complete SV40 DNA replication with purified replication factors. The Journal of biological chemistry 269, 10923–10934

7. Kunkel, T. A., and Burgers, P. M. (2008) Dividing the workload at a eukaryotic replication fork. Trends in cell biology 18, 521–527

8. MacNeill, S. A., Baldacci, G., Burgers, P. M., and Hubscher, U. (2001) A unified nomenclature for the subunits of eukaryotic DNA polymerase delta. Trends in biochemical sciences 26, 16–17

9. Miyabe, I., Kunkel, T. A., and Carr, A. M. (2011) The major roles of DNA polymerases epsilon and delta at the eukaryotic replication fork are evolutionarily conserved. PLoS genetics 7, e1002407

10. Zuo, S., Bermudez, V., Zhang, G., Kelman, Z., and Hurwitz, J. (2000) Structure and activity associated with multiple forms of Schizosaccharomyces pombe DNA polymerase delta. The Journal of biological chemistry 275, 5153–5162

11. Zhou, Y., Meng, X., Zhang, S., Lee, E. Y., and Lee, M. Y. (2012) Characterization of human DNA polymerase delta and its subassemblies reconstituted by expression in the MultiBac system. PloS one 7, e39156

12. Johansson, E., Garg, P., and Burgers, P. M. (2004) The Pol32 subunit of DNA polymerase delta contains separable domains for processive replication and proliferating cell nuclear antigen (PCNA) binding. The Journal of biological chemistry 279, 1907–1915

13. Bermudez, V. P., MacNeill, S. A., Tappin, I., and Hurwitz, J. (2002) The influence of the Cdc27 subunit on the properties of the Schizosaccharomyces pombe DNA polymerase delta. The Journal of biological chemistry 277, 36853–36862

14. Li, H., Xie, B., Zhou, Y., Rahmeh, A., Trusa, S., Zhang, S., Gao, Y., Lee, E. Y., and Lee, M. Y. (2006) Functional roles of p12, the fourth subunit of human DNA polymerase delta. The Journal of biological chemistry 281, 14748–14755

15. Lee, M. Y., Zhang, S., Lin, S. H., Chea, J., Wang, X., LeRoy, C., Wong, A., Zhang, Z., and Lee, E. Y. (2012) Regulation of human DNA polymerase delta in the cellular responses to DNA damage. Environmental and molecular mutagenesis 53, 683–698

16. Lee, M. Y., Zhang, S., Lin, S. H., Wang, X., Darzynkiewicz, Z., Zhang, Z., and Lee, E. Y. (2014) The tail that wags the dog: p12, the smallest subunit of DNA polymerase delta, is degraded by ubiquitin ligases in response to DNA damage and during cell cycle progression. Cell cycle 13, 23–31

17. Meng, X., Zhou, Y., Lee, E. Y., Lee, M. Y., and Frick, D. N. (2010) The p12 subunit of human polymerase delta modulates the rate and fidelity of DNA synthesis. Biochemistry 49, 3545–3554

18. Krishna, T. S., Kong, X. P., Gary, S., Burgers, P. M., and Kuriyan, J. (1994) Crystal structure of the eukaryotic DNA polymerase processivity factor PCNA. Cell 79, 1233–1243

19. Wang, Y., Zhang, Q., Chen, H., Li, X., Mai, W., Chen, K., Zhang, S., Lee, E. Y., Lee, M. Y., and Zhou, Y. (2011) P50, the small subunit of DNA polymerase delta, is required for mediation of the interaction of polymerase delta subassemblies with PCNA. PloS one 6, e27092

20. Lu, X., Tan, C. K., Zhou, J. Q., You, M., Carastro, L. M., Downey, K. M., and So, A. G. (2002) Direct interaction of proliferating cell nuclear antigen with the small subunit of DNA polymerase delta. The Journal of biological chemistry 277, 24340–24345

21. Zhang, P., Mo, J. Y., Perez, A., Leon, A., Liu, L., Mazloum, N., Xu, H., and Lee, M. Y. (1999) Direct interaction of proliferating cell nuclear antigen with the p125 catalytic subunit of mammalian DNA polymerase delta. The Journal of biological chemistry 274, 26647–26653

22. Terai, K., Shibata, E., Abbas, T., and Dutta, A. (2013) Degradation of p12 subunit by CRL4Cdt2 E3 ligase inhibits fork progression after DNA damage. The Journal of biological chemistry 288, 30509–30514

23. Rahmeh, A. A., Zhou, Y., Xie, B., Li, H., Lee, E. Y., and Lee, M. Y. (2012) Phosphorylation of the p68 subunit of Pol delta acts as a molecular switch to regulate its interaction with PCNA. Biochemistry 51, 416–424

24. Fazlieva, R., Spittle, C. S., Morrissey, D., Hayashi, H., Yan, H., and Matsumoto, Y. (2009) Proofreading exonuclease activity of human DNA polymerase delta and its effects on lesion-bypass DNA synthesis. Nucleic acids research 37, 2854–2866

25. Podust, V. N., Chang, L. S., Ott, R., Dianov, G. L., and Fanning, E. (2002) Reconstitution of human DNA polymerase delta using recombinant baculoviruses: the p12 subunit potentiates DNA polymerizing activity of the four-subunit enzyme. The Journal of biological chemistry 277, 3894–3901

26. Essers, J., Theil, A. F., Baldeyron, C., van Cappellen, W. A., Houtsmuller, A. B., Kanaar, R., and Vermeulen, W. (2005) Nuclear dynamics of PCNA in DNA replication and repair. Molecular and cellular biology 25, 9350–9359

27. Ducoux, M., Urbach, S., Baldacci, G., Hubscher, U., Koundrioukoff, S., Christensen, J., and Hughes, P. (2001) Mediation of proliferating cell nuclear antigen (PCNA)-dependent DNA replication through a conserved p21(Cip1)-like PCNA-binding motif present in the third subunit of human DNA polymerase delta. The Journal of biological chemistry 276, 49258–49266

28. Pohler, J. R., Otterlei, M., and Warbrick, E. (2005) An in vivo analysis of the localisation and interactions of human p66 DNA polymerase delta subunit. BMC molecular biology 6, 17

29. Zerangue, N., Schwappach, B., Jan, Y. N., and Jan, L. Y. (1999) A new ER trafficking signal regulates the subunit stoichiometry of plasma membrane K(ATP) channels. Neuron 22, 537–548

30. Joiner, W. J., Wang, L. Y., Tang, M. D., and Kaczmarek, L. K. (1997) hSK4, a member of a novel subfamily of calcium-activated potassium channels. Proceedings of the National Academy of Sciences of the United States of America 94, 11013–11018

31. Jones, H. M., Bailey, M. A., Baty, C. J., Macgregor, G. G., Syme, C. A., Hamilton, K. L., and Devor, D. C. (2007) An NH2-terminal multi-basic RKR motif is required for the ATP-dependent regulation of hIK1. Channels (Austin) 1, 80–91

32. Fagerlund, R., Melen, K., Kinnunen, L., and Julkunen, I. (2002) Arginine/lysine-rich nuclear localization signals mediate interactions between dimeric STATs and importin alpha 5. The Journal of biological chemistry 277, 30072–30078

33. Collins, P. E., Kiely, P. A., and Carmody, R. J. (2014) Inhibition of transcription by B cell Leukemia 3 (Bcl-3) protein requires interaction with nuclear factor kappaB (NF-kappaB) p50. The Journal of biological chemistry 289, 7059–7067

34. van Hennik, P. B., ten Klooster, J. P., Halstead, J. R., Voermans, C., Anthony, E. C., Divecha, N., and Hordijk, P. L. (2003) The C-terminal domain of Rac1 contains two motifs that control targeting and signaling specificity. The Journal of biological chemistry 278, 39166–39175

35. Wang, L., Zhang, K., Wu, L., Liu, S., Zhang, H., Zhou, Q., Tong, L., Sun, F., and Fan, Z. (2012) Structural insights into the substrate specificity of human granzyme H: the functional roles of a novel RKR motif. Journal of immunology 188, 765–773

36. Naryzhny, S. N., Desouza, L. V., Siu, K. W., and Lee, H. (2006) Characterization of the human proliferating cell nuclear antigen physico-chemical properties: aspects of double trimer stability. Biochemistry and cell biology = Biochimie et biologie cellulaire 84, 669–676

37. Manohar, K., and Acharya, N. (2015) Characterization of proliferating cell nuclear antigen (PCNA) from pathogenic yeast Candida albicans and its functional analyses in S. Cerevisiae. BMC microbiology 15, 257

38. Xie, B., Mazloum, N., Liu, L., Rahmeh, A., Li, H., and Lee, M. Y. (2002) Reconstitution and characterization of the human DNA polymerase delta four-subunit holoenzyme. Biochemistry 41, 13133–13142

39. Acharya, N., Haracska, L., Johnson, R. E., Unk, I., Prakash, S., and Prakash, L. (2005) Complex formation of yeast Rev1 and Rev7 proteins: a novel role for the polymerase-associated domain. Molecular and cellular biology 25, 9734–9740

40. Burgers, P. M. (1998) Eukaryotic DNA polymerases in DNA replication and DNA repair. Chromosoma 107, 218–227

41. Hindges, R., and Hubscher, U. (1997) DNA polymerase delta, an essential enzyme for DNA transactions. Biological chemistry 378, 345–362

42. Albertson, T. M., Ogawa, M., Bugni, J. M., Hays, L. E., Chen, Y., Wang, Y., Treuting, P. M., Heddle, J. A., Goldsby, R. E., and Preston, B. D. (2009) DNA polymerase epsilon and delta proofreading suppress discrete mutator and cancer phenotypes in mice. Proceedings of the National Academy of Sciences of the United States of America 106, 17101–17104

43. Flohr, T., Dai, J. C., Buttner, J., Popanda, O., Hagmuller, E., and Thielmann, H. W. (1999) Detection of mutations in the DNA polymerase delta gene of human sporadic colorectal cancers and colon cancer cell lines. International journal of cancer 80, 919–929

44. Goldsby, R. E., Hays, L. E., Chen, X., Olmsted, E. A., Slayton, W. B., Spangrude, G. J., and Preston, B. D. (2002) High incidence of epithelial cancers in mice deficient for DNA polymerase delta proofreading. Proceedings of the National Academy of Sciences of the United States of America 99, 15560–15565

45. Lee, M., Wang, X., Zhang, S., Zhang, Z., and Lee, E. Y. C. (2017) Regulation and Modulation of Human DNA Polymerase delta Activity and Function. Genes 8

46. Zhang, Q., Zhang, Q., Chen, H., Chen, Y., and Zhou, Y. (2016) Multiple Forms of Human DNA Polymerase Delta Sub-Assembling in Cellular DNA Transactions. Current protein & peptide science 17, 746–755

47. Zhou, Y., Chen, H., Li, X., Wang, Y., Chen, K., Zhang, S., Meng, X., Lee, E. Y., and Lee, M. Y. (2011) Production of recombinant human DNA polymerase delta in a Bombyx mori bioreactor. PloS one 6, e22224

48. Rizzo, A. A., Vassel, F. M., Chatterjee, N., D’Souza, S., Li, Y., Hao, B., Hemann, M. T., Walker, G. C., and Korzhnev, D. M. (2018) Rev7 dimerization is important for assembly and function of the Rev1/Polzeta translesion synthesis complex. Proceedings of the National Academy of Sciences of the United States of America 115, E8191–E8200

49. Bruning, J. B., and Shamoo, Y. (2004) Structural and thermodynamic analysis of human PCNA with peptides derived from DNA polymerase-delta p66 subunit and flap endonuclease-1. Structure 12, 2209–2219

50. Lemmens, L., Urbach, S., Prudent, R., Cochet, C., Baldacci, G., and Hughes, P. (2008) Phosphorylation of the C subunit (p66) of human DNA polymerase delta. Biochemical and biophysical research communications 367, 264–270

51. Baranovskiy, A. G., Babayeva, N. D., Liston, V. G., Rogozin, I. B., Koonin, E. V., Pavlov, Y. I., Vassylyev, D. G., and Tahirov, T. H. (2008) X-ray structure of the complex of regulatory subunits of human DNA polymerase delta. Cell cycle 7, 3026–3036

52. Chea, J., Zhang, S., Zhao, H., Zhang, Z., Lee, E. Y., Darzynkiewicz, Z., and Lee, M. Y. (2012) Spatiotemporal recruitment of human DNA polymerase delta to sites of UV damage. Cell cycle 11, 2885–2895

53. Fan, X., Zhang, Q., You, C., Qian, Y., Gao, J., Liu, P., Chen, H., Song, H., Chen, Y., Chen, K., and Zhou, Y. (2014) Proteolysis of the human DNA polymerase delta smallest subunit p12 by mu-calpain in calcium-triggered apoptotic HeLa cells. PloS one 9, e93642

54. Eissenberg, J. C., Ayyagari, R., Gomes, X. V., and Burgers, P. M. (1997) Mutations in yeast proliferating cell nuclear antigen define distinct sites for interaction with DNA polymerase delta and DNA polymerase epsilon. Molecular and cellular biology 17, 6367–6378

55. Gomes, X. V., and Burgers, P. M. (2000) Two modes of FEN1 binding to PCNA regulated by DNA. The EMBO journal 19, 3811–3821

56. Murga, M., Lecona, E., Kamileri, I., Diaz, M., Lugli, N., Sotiriou, S. K., Anton, M. E., Mendez, J., Halazonetis, T. D., and Fernandez-Capetillo, O. (2016) POLD3 Is Haploinsufficient for DNA Replication in Mice. Molecular cell 63, 877–883

57. Schmitt, M. W., Matsumoto, Y., and Loeb, L. A. (2009) High fidelity and lesion bypass capability of human DNA polymerase delta. Biochimie 91, 1163–1172

