## supplementary fig. and legend for "Human DNA polymerase delta is a pentameric holoenzyme with dimeric p12 subunit: Implications in enzyme architecture and PCNA interaction"

*Correspondence to:

**Supplementary Fig. 1:** GST-affinity purification of hPolδ holoenzyme was carried out by using bacterial expression constructs harboring GST-p125 and a triple expressing plasmid containing p12, p50 and p68. Both p125 and p12 possess protease site. Cleaved p12 retains FLAG–tag and thus it migrates ~22Kda position. * Bacterial chaperon and other non-specific proteins. Stochiometry of FLAG-p12 is higher than any other subunit.  Indicated bands were also authenticated by western blot analysis.

**Supplementary Fig. 2:** p12 RKR (1-MGRKRLITDSYPVK-14) and PIP (92-GDPRFQCSLWHLYPL-106) motif harboring domains were used for peptide structure prediction by using PEP-FOLD3 server (http://bioserv.rpbs.univ-paris-diderot.fr/services/PEP-FOLD3/). The models were validated by SAVES and Ramachandran plot. RKR-motif forms a stable α-helix, whereas PIP motif forms a $3_{10}$ helix.

**Supplementary Fig. 3:** p12 proteins were purified to near homogeneity from bacterial cells by using GST-affinity column chromatography, GST-tag was cleaved off by PreScission protease, and analyzed by SDS containing polyacrylamide gel electrophoresis (PAGE). Lane 1, MW; Lane 2, CA; Lane 3 wild type;  Lane 4 R3A, K4A, R5A and Lane 5 L104A, Y105A p12 proteins.

**Supplementary table- 1:** List of primers used for various orfs amplification with underlined restriction enzymes.

| Primer | Primer sequence (5'----3') | Name of the ORF |
|---|---|---|
| **NAP 239** | ccggggatccacatatgttcgaggcgcgcc | hPCNA |
| **NAP 240** | ccggggatccctaagatccttcttcatc | hPCNA |
| **NAP 251** | ccggggatccgtatgttcgaggcgcgcc | hPCNA |
| **NAP 300** | ctacacagctgtactcctgttctggagctccagcttgttcaacatc | hPCNA |
| **NAP 304** | gatgttgaacaagctggagctccagaacaggagtacagctgtgtag | hPCNA |
| **NAP 305** | ggccggatccctaagatccttcttcatcctcgaggcggcagccaag | hPCNA |
| **NAP 254** | ggccggatcctcaggggcccagccccagg | p50 |
| **NAP 255** | ggcgaattcatgttttctgagcaggctgc | p50 |
| **NAP 257** | ggccggatccttatttcctctggaagaagcc | p68 |
| **NAP 258** | ggcgaattcatggcggaccagctttatctgg | p68 |
| **NAP 260** | ccggggatccacatatgggccggaagcggctc | p12 |
| **NAP 261** | ggccggatcctcataggggatagagatgcc | p12 |
| **NAP 262** | ggcgaattcatgggccggaagcggctc | p12 |
| **NAP 265** | ggccggatcctcataggggagcggcatgccagagactg | p12 |
| **NAP 362** | ccggggatccgtatgggcgccgctgcactcatcactgattcc | p12 |
| **NAP 373** | ggccggatccacatatgggcgccgctgcactcatcactg | p12 |
| **NAP 252** | ggcgaattcatggatggcaagcggcgg | p125 |
| **NAP 248** | ggccggatcctcaccaggcctcaggtccagggggtcc | p125 |
| **NAP 361** | ggccggatccctagcctggaatggtttgaag | Cdm1 |
| **NAP 451** | ccggggatccacatatgaagaagcgcac | Cdm1 |
| **NAP 452** | ggccggatccacatatgactactcaagcgaaaaaatcaggg | Cdm1 |
| **NAP 448** | cggcgaattctatgggccggaagcgg | p12 |
| **NAP 450** | ccggggatccggtaggggatgatagagatg | p12 |
| **NAP 444** | ccgggaattctatgctagaaaacaatgc | Polθ |
| **NAP 151** | ccgggtcgacggatccttacacatcaaagtcctttagctctcccc | Polθ |

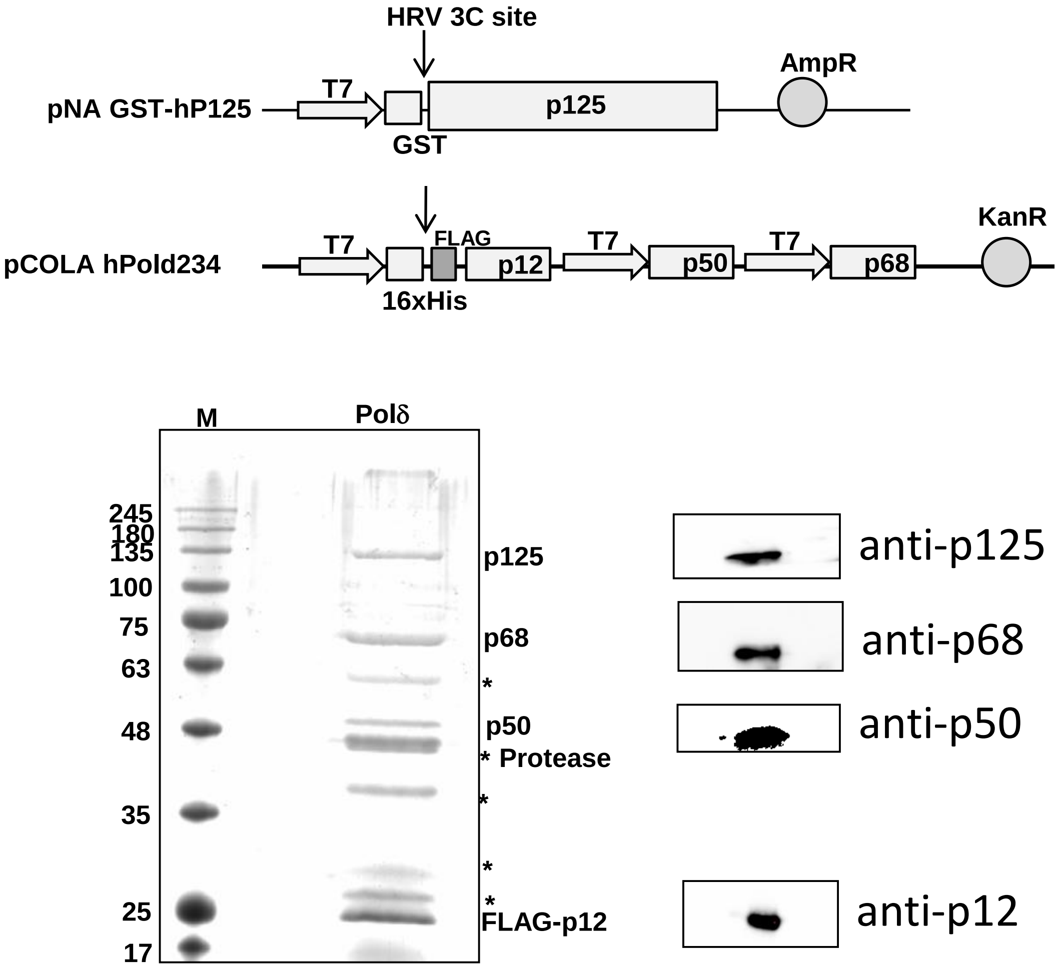


**Supp. Figure. 1**

**A.**

1-MGRKRLITDSYPVVKRREG-19 89-KTHPGDPRFQCSLWHLYPL-107

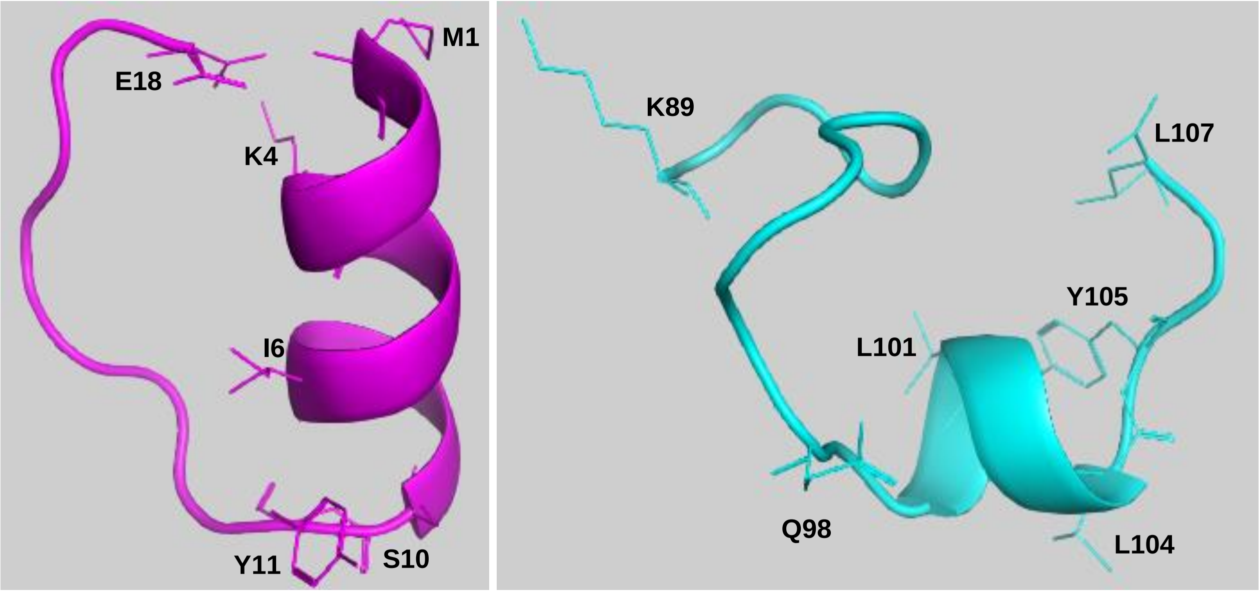


**B.**

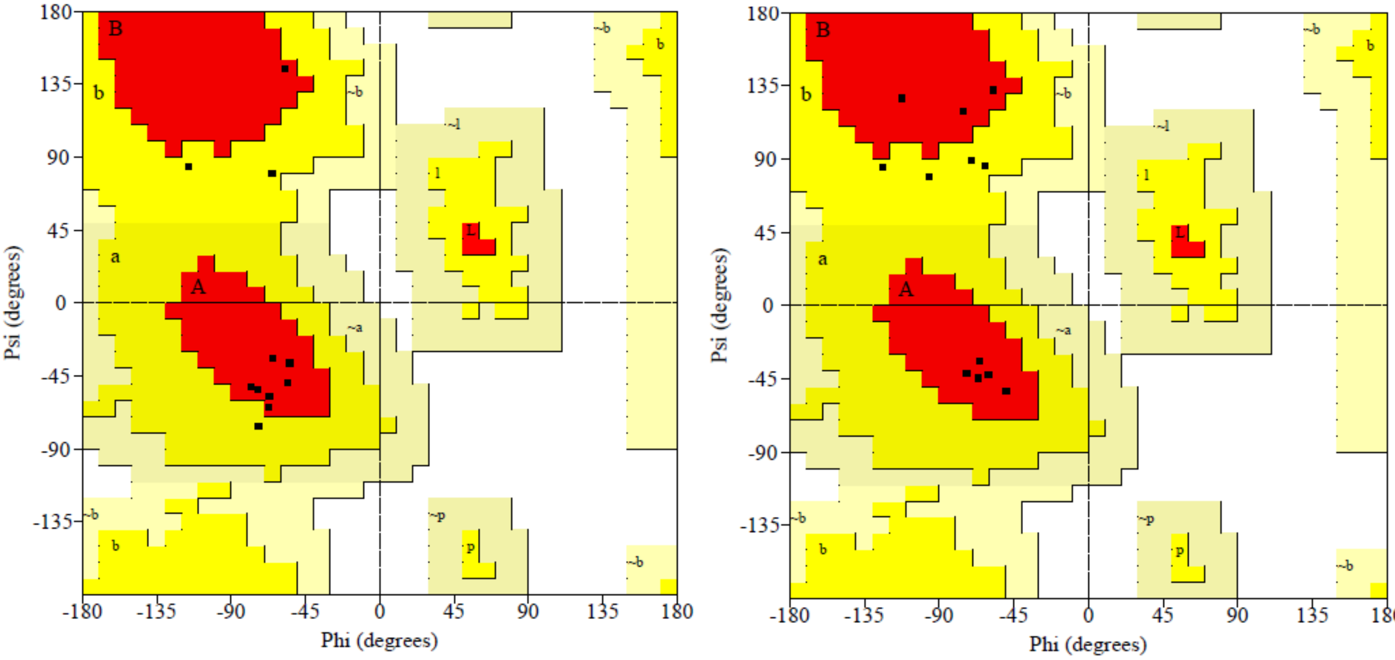


**Supp. Figure. 2**

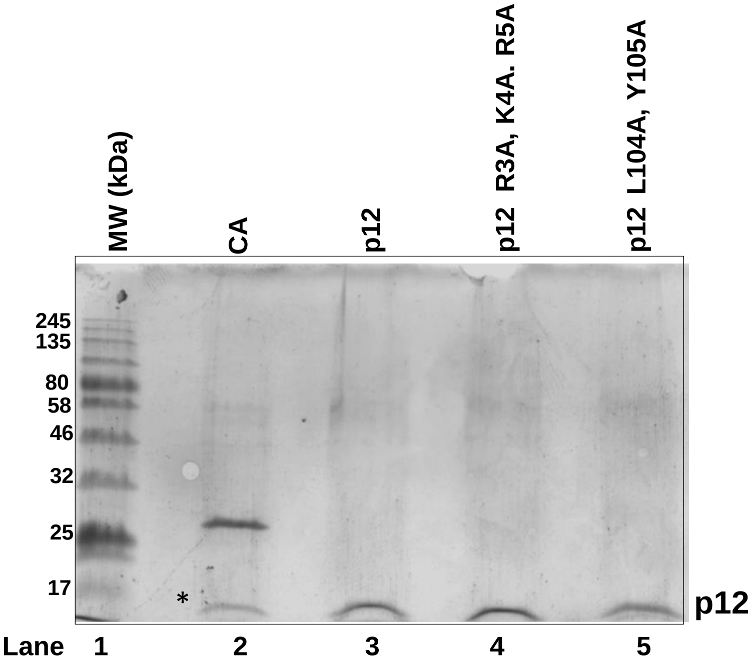


**Supp. Figure. 3**
